# High-Efficiency Pharmacogenetic Ablation of Oligodendrocyte Progenitor Cells in the Adult Mouse CNS

**DOI:** 10.1101/2021.05.13.443012

**Authors:** Yao Lulu Xing, Jasmine Poh, Bernard H.A. Chuang, Kaveh Moradi, Stanislaw Mitew, William D. Richardson, Trevor J. Kilpatrick, Yasuyuki Osanai, Tobias D. Merson

**Affiliations:** Australian Regenerative Medicine Institute, Monash University, Clayton, Victoria, Australia; Florey Institute of Neuroscience and Mental Health, Parkville, Victoria, Australia; Wolfson Institute for Biomedical Research, University College London, Gower Street, London, United Kingdom; Current address: Division of Histology and Cell Biology, Department of Anatomy, Jichi Medical University, Tochigi, Japan; Current address: National Institute of Mental Health, National Institutes of Health, Bethesda, MD, USA; Corresponding author and lead contact.

## Abstract

Approaches to investigate adult oligodendrocyte progenitor cells (OPCs) by targeted cell ablation in the rodent central nervous system have been limited by methodological challenges resulting in only partial and transient OPC depletion. We have developed a novel pharmacogenetic model of conditional OPC ablation, eliminating 98.6% of all OPCs throughout the brain. By combining recombinase-based transgenic and viral strategies for targeting OPCs and ventricular-subventricular zone (V-SVZ)-derived neural precursor cells (NPCs), we found new PDGFRA-expressing cells born in the V-SVZ repopulated the OPC-deficient brain starting 12 days after OPC ablation. Our data reveal that OPC depletion induces V-SVZ-derived NPCs to generate vast numbers of PDGFRA^+^ NG2^+^ cells with the capacity to migrate and proliferate extensively throughout the dorsal anterior forebrain. Further application of this novel approach to ablate OPCs will advance knowledge of the function of both OPCs and oligodendrogenic NPCs in health and disease.

## Introduction

Oligodendrocyte progenitor cells (OPCs), also known as NG2-glia, are the principal mitotic cell type in the adult mammalian central nervous system (CNS). OPCs are known primarily for generating myelin-forming oligodendrocytes (OLs) during postnatal development and adulthood (Kang et al., 2010; Young et al., 2013). While OPCs are distributed more or less uniformly throughout the CNS, including in brain regions where relatively little myelination occurs, there is evidence for OPC heterogeneity between brain regions (Chittajallu et al., 2004; Spitzer et al., 2019), which has raised the prospect that they could possess additional functions beyond oligodendrogenesis.

To investigate the function of OPCs within the adult CNS, several groups have developed different strategies to ablate OPCs selectively. However, these experimental approaches, including X-irradiation, laser-mediated ablation, genetically-induced cell ablation, or the use of anti-mitotic drugs, have their specific caveats that have limited the extent to which it has been possible to achieve long-term depletion of OPCs in adult mice (Birey et al., 2015; Dang et al., 2019; Hughes et al., 2013; Irvine and Blakemore, 2007; Nakano et al., 2017; Robins et al., 2013). Existing approaches have enabled only partial and transient OPC ablation throughout the CNS due to incomplete targeting of the OPC population and rapid repopulation by nearby non-ablated OPCs. As a result, it has not yet been possible to explore the functional consequences of long-term OPC ablation.

Designing a genetic approach to completely and selectively ablate OPCs in the adult mouse CNS requires careful consideration of the promoter(s) used to target the OPC population. Parenchymal OPCs are defined by their expression of chondroitin sulfate proteoglycan 4/neuron-glial antigen 2 (CSPG4/NG2) (Nishiyama et al., 2009) and platelet-derived growth factor receptor alpha (PDGFRA) (Rivers et al., 2008). Although the *Cspg4* promoter has been used to control transgene expression for studies of parenchymal OPC fate, behavior, and function (Birey *et al*., 2015; Chang et al., 2012; Kang *et al*., 2010; Ziskin et al., 2007), NG2 is also expressed by pericytes (Marques et al., 2016; Nakano *et al*., 2017; Robins *et al*., 2013; Vanlandewijck et al., 2018) and by some microglial cells after injury (Zhu et al., 2016). Similarly, PDGFRA is not an exclusive marker of OPCs; PDGFRA is also expressed by vascular and leptomeningeal cells (VLMCs, also known as perivascular fibroblasts) (Marques *et al*., 2016; Vanlandewijck *et al*., 2018) and by choroid plexus epithelial cells (Kang *et al*., 2010; Pringle et al., 1992; Schatteman et al., 1992). Therefore, using either the *Cspg4* or *Pdgfra* promoter alone cannot direct the expression of a suicide gene exclusively to OPCs. To precisely target OPCs without affecting other neural cell types, we have generated a novel transgenic mouse model in which the expression of an inducible suicide gene is controlled by two different promoters, namely the *Pdgfra* and *Sox10* promoters, whose overlapping transcriptional activity is restricted to OPCs in the postnatal CNS.

In addition to the specificity of genetic targeting, the method to conditionally ablate OPCs must also be highly efficient, targeting most, if not all, OPCs. High efficiency is critical for overcoming the proliferative response of non-ablated OPCs that follows incomplete OPC ablation, which has been demonstrated to result in swift regeneration of the OPC population (Birey *et al*., 2015; Hughes *et al*., 2013; Robins *et al*., 2013). Moreover, the method should be amenable to precise temporal control and have minimal effect on the animal’s overall health. To date, no strategy for ablating the entire OPC population has been described that meets all of these requirements.

We have developed a pharmacogenetic approach to ablate OPCs in the adult mouse CNS that overcomes many limitations of previous approaches. The method involves the inducible and conditional expression of diphtheria toxin A (DTA) in adult OPCs followed by delivery of an anti-mitotic agent into the CNS to ablate dividing OPCs that escape genetic targeting. We demonstrate that this approach provides highly efficient and selective ablation of OPCs across the entire adult mouse brain, resulting in the loss of 98.6 ± 0.4% of all OPCs.

Using this pharmacogenetic model of conditional OPC ablation also uncovered the remarkable compensatory regenerative response of the ventricular-subventricular zone (V-SVZ). Beginning 12 days after widespread OPC ablation, Nestin-expressing neural precursor cells (NPCs) residing in the V-SVZ generated new PDGFRA^+^ NG2^+^ cells whose morphology resembled those of OPCs. The newly generated PDGFRA^+^ cells migrated extensively throughout the dorsal anterior forebrain, re-establishing the broad distribution of OPCs in this region of the brain. Co-ablation of both OPCs and oligodendrogenic NPCs prevented the regeneration of PDGFRA^+^ cells in the cerebrum. Collectively, our mouse model of conditional OPC ablation provides a valuable experimental tool for future studies aimed at better understanding the functions of OPCs and for further exploring the oligodendrogenic potential of NPCs residing in the V-SVZ.

## Results

### DTA-mediated ablation of OPCs induced rapid regeneration of the OPC population

To specifically ablate OPCs in the adult mouse CNS, we used an intersectional genetic approach to direct the inducible expression of a suicide gene in cells expressing both PDGFRA and SOX10. This was achieved by crossing two transgenic mouse lines, the *Pdgfrα-CreER^T2^* line (Rivers *et al*., 2008) and the *Sox10-lox-GFP-STOP-lox-DTA* (*Sox10-DTA*) line (Kessaris et al., 2006), to enable DTA expression in adult OPCs upon delivery of tamoxifen (TAM) (**Figure 1A**). Since the SOX10 transcription factor is expressed exclusively by cells of the oligodendroglial lineage in the postnatal CNS (Lu et al., 2000; Xu et al., 2000), this ensures that DTA expression is restricted to OPCs and is not induced in either VLMCs or choroid plexus epithelial cells, both of which express PDGFRA but not SOX10. Similarly, DTA expression is not expected to target Schwann cells in the peripheral nervous system since the majority of Schwann cells express SOX10 but not PDGFRA (Britsch et al., 2001; Eccleston et al., 1993). Moreover, it has been demonstrated conclusively that the *Pdgfrα-CreER^T2^* line used in our study does not target Schwann cells (Zawadzka et al., 2010).

**Figure 1.**
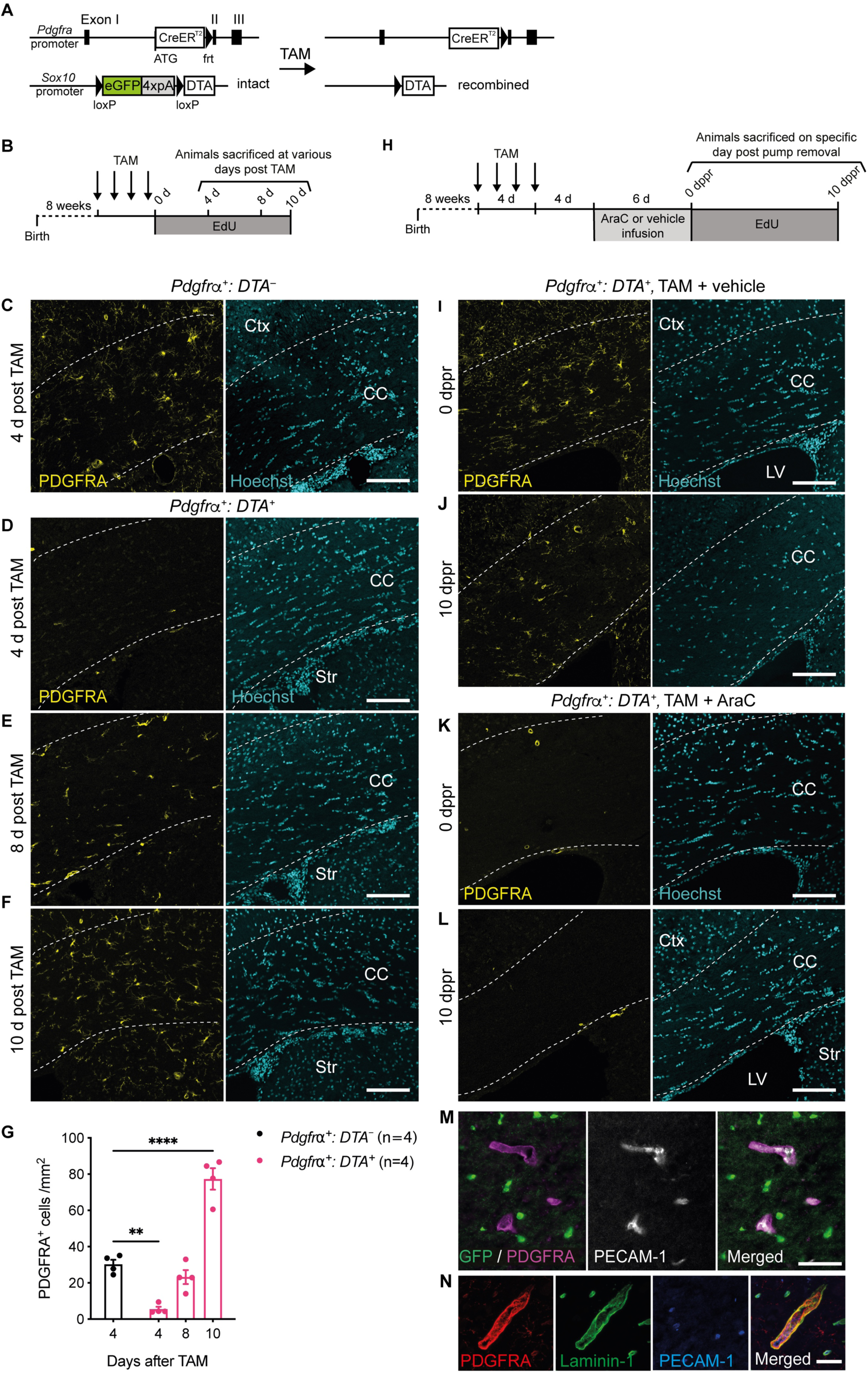
TAM administration followed by intracisternal infusion of AraC resulted in the ablation of almost all OPCs in the brain. **A**, Schematic of the transgenic alleles in *Pdgfrα^+^: DTA^+^* mice. **B,** Timeline for the assessment of DTA-mediated OPC ablation after TAM administration. **C**, Coronal brain sections of a TAM - administered *Pdgfrα ^+^: DTA^−^* mouse immunolabeled with antibodies against GFP and PDGFRA revealed abundant OPCs in the corpus callosum (CC). **D**, Lack of PDGFRA^+^ OPCs in the corpus callosum of a *Pdgfrα ^+^: DTA^+^*mouse sacrificed 4 days after TAM administration. **E,F**, Regeneration of PDGFRA^+^ OPCs at days 8 and 10 after final TAM administration. **G**, Density of PDGFRA^+^ OPCs in the corpus callosum of *Pdgfrα^+^: DTA^+^* mice at 4, 8 and 10 days after TAM administration. **H**, Timeline for the assessment of DTA-mediated OPC ablation following TAM + AraC administration. **I,J,** OPC repopulation in a vehicle-infused transgenic mouse assessed at the end of infusion (**I**) or 10 days later (**J**). **K,L,** OPC repopulation was not observed in the corpus callosum of an AraC-infused mouse at either 0 or 10 dppr. **M,N,** Residual PDGFRA-expressing cells in an AraC-infused mouse collected at 10 dppr were identified as perivascular fibroblast-like cells based on their close association with PECAM-1-labelled endothelial cells (**M**) and colocalization of PDGFRA and Laminin-1 (**N**). CC, corpus callosum; Ctx, cerebral cortex; LV, lateral ventricle; TAM, tamoxifen. Data represent mean ± SEM. Statistical analysis: Two-way ANOVA with Dunnett’s multiple comparison tests, \*\**p*<0.01, \*\*\*\**p*<0.0001. Scale bars, 100 μm (**C-F**, **I-L**), 50 μm **(M**) and 30 µm (**N**).

TAM was administered to 8-week-old *Pdgfrα-CreER^T2+/+^: Sox10-DTA^+/−^* mice (hereafter denoted *Pdgfrα^+^: DTA^+^*), as well as to *Pdgfrα-CreER^T2+/+^: Sox10-DTA^−/−^* littermates lacking the *Sox10-DTA* allele (denoted *Pdgfrα^+^: DTA^−^*), which served as non-ablated controls (**Figure 1B**). Immunohistochemistry on the brains of non-ablated *Pdgfrα^+^: DTA^−^* controls sacrificed 4 days after the final dose of TAM revealed abundant PDGFRA^+^ OPCs throughout the brain, including the corpus callosum (**Figure 1C**). By contrast, TAM administered *Pdgfrα^+^: DTA^+^* mice assessed at the same time-point had very few PDGFRA^+^ OPCs in the corpus callosum (**Figure 1D**), consistent with the notion that Cre-mediated induction of DTA resulted in OPC ablation. In these OPC-deficient mice, *Sox10* promoter-driven GFP expression was restricted to SOX10^+^ CC1^+^ cells (**Figure S1A,B**), indicating that Cre-mediated recombination of the *Sox10-DTA* allele targeted OPCs but not mature OLs.

Although *Pdgfrα^+^: DTA^+^* mice exhibited marked OPC depletion 4 days after TAM administration, OPC density in the corpus callosum returned to control levels by day 8 post TAM and increased further over the subsequent 2 days (**Figure 1E-G**). The marked increase in OPC density observed 10 days after TAM delivery suggests that OPCs exhibit robust proliferation as a compensatory response to acute ablation. Consistent with this observation, most OPCs present after 10 days were newly-generated, as demonstrated by the significant proportion of PDGFRA^+^ cells that had incorporated 5-ethynyl-2ʹ-deoxyuridine (EdU) which was provided continuously in the drinking water after the last TAM administration (**Figure S1C,D**). Most OPCs present 4 days after TAM administration expressed GFP (**Figure S1D**), suggesting that the surviving OPCs were principally those in which the *Sox10-DTA* allele had not recombined. Some EdU^+^ OPCs did not express GFP (**Figure S1D**), which most likely reflects low transcriptional activity of the *Sox10* promoter that directs GFP expression. Supporting this idea, not all SOX10^+^ oligodendroglia in *Pdgfrα^+^: DTA^+^* brains examined 4 days post TAM expressed GFP (**Figure S1A**). These data demonstrate that the vast majority of OPCs in *Pdgfrα^+^: DTA^+^* mice were depleted after TAM administration. However, residual non-recombined OPCs exhibited a robust proliferative response to OPC ablation, resulting in restoration of OPCs to similar or higher density than non-ablated controls within 8-10 days after TAM administration.

### Intracisternal infusion of AraC after TAM administration prevented rapid OPC regeneration

Given that incomplete OPC ablation triggered non-recombined OPCs that had escaped DTA-mediated apoptosis to proliferate and repopulate the CNS, we introduced a second intervention designed to kill these rapidly dividing OPCs. Specifically, after TAM administration, the anti-mitotic drug cytosine-β-D-arabinofuranoside (AraC) was infused into the cisterna magna of *Pdgfrα^+^: DTA^+^* mice to deplete all residual proliferating OPCs. We elected to administer AraC directly into the cerebrospinal fluid rather than providing additional doses of TAM, given that we have noted toxicity when administering TAM for more than 4 consecutive days. Osmotic minipumps were implanted on day 4 after TAM administration and removed on day 10 (**Figure 1H**), to provide 6 days of continuous AraC infusion during the period of marked OPC proliferation. Vehicle-only controls received artificial cerebrospinal fluid (aCSF) without AraC. Mice were humanely sacrificed either immediately after removal of the osmotic minipump or 10 days later (**Figure 1H**).

TAM-administered *Pdgfrα^+^: DTA^+^* mice examined after 6 days of vehicle infusion, denoted as 0 days post pump removal (dppr), had numerous PDGFRA^+^ OPCs in the corpus callosum (**Figure 1I**), similar in density to that observed in *Pdgfrα^+^: DTA^+^* mice administered TAM alone and assessed 10 days later (**Figure 1F**). By contrast, almost no PDGFRA^+^ OPCs could be identified in the corpus callosum of TAM-administered *Pdgfrα^+^: DTA^+^* mice sacrificed immediately after AraC infusion (**Figure 1K**). OPC depletion was sustained for at least 10 days after removal of the osmotic minipump used to deliver AraC (**Figure 1L**). By contrast, AraC delivery to wild-type mice did not result in a decrement in OPC density (**Figure S1E**), consistent with the low proliferation rate of OPCs under basal conditions (Gonsalvez et al., 2019; Marques *et al*., 2016) and known homeostatic control mechanisms that maintain OPC density in equilibrium (Hughes *et al*., 2013). While PDGFRA^+^ OPCs were almost completely absent in TAM + AraC administered *Pdgfrα^+^: DTA^+^* mice, vascular-associated PDGFRA^+^ GFP^−^ cells surrounding PECAM-1^+^ endothelial cells in the brain parenchyma remained intact (**Figure 1M**). These vascular-associated PDGFRA^+^ GFP^−^ cells in OPC-ablated mice did not exhibit typical ramified OPC morphology. We identified these vascular-associated PDGFRA^+^ cells as laminin-1^+^ perivascular fibroblast-like cells that are closely associated with but distinct from vascular-associated NG2^+^ PDGFRB^+^ pericytes (**Figure 1N, Figure S2A-D**), consistent with the recent description of these cells elsewhere (Vanlandewijck *et al*., 2018).

To explore the extent of OPC ablation in greater detail, we generated *Pdgfrα-CreER^T2+/–^: Ai14-tdTomato^+/−^: Sox10-DTA^+/−^* mice (hereafter denoted *Pdgfrα^+^: tdT^+^: DTA^+^*), to enable simultaneous genetic fate-mapping and ablation of OPCs. TAM was administered to 8-week-old *Pdgfrα^+^: tdT^+^: DTA^+^* mice to induce expression of both tdTomato and DTA from the *Ai14 tdTomato* and *Sox10-DTA* recombined alleles, respectively (**Figure 2A**). Four days after the last dose of TAM, mice received a 6-day intracisternal infusion of AraC, and their brains were collected immediately after infusion to analyze the extent of OPC depletion. *Pdgfrα^+^: tdT^+^: DTA^−^* littermates administered TAM and infused with vehicle alone served as non-ablated controls and allowed us to determine the specificity and efficiency of tdTomato expression within the intact OPC population.

**Figure 2.**
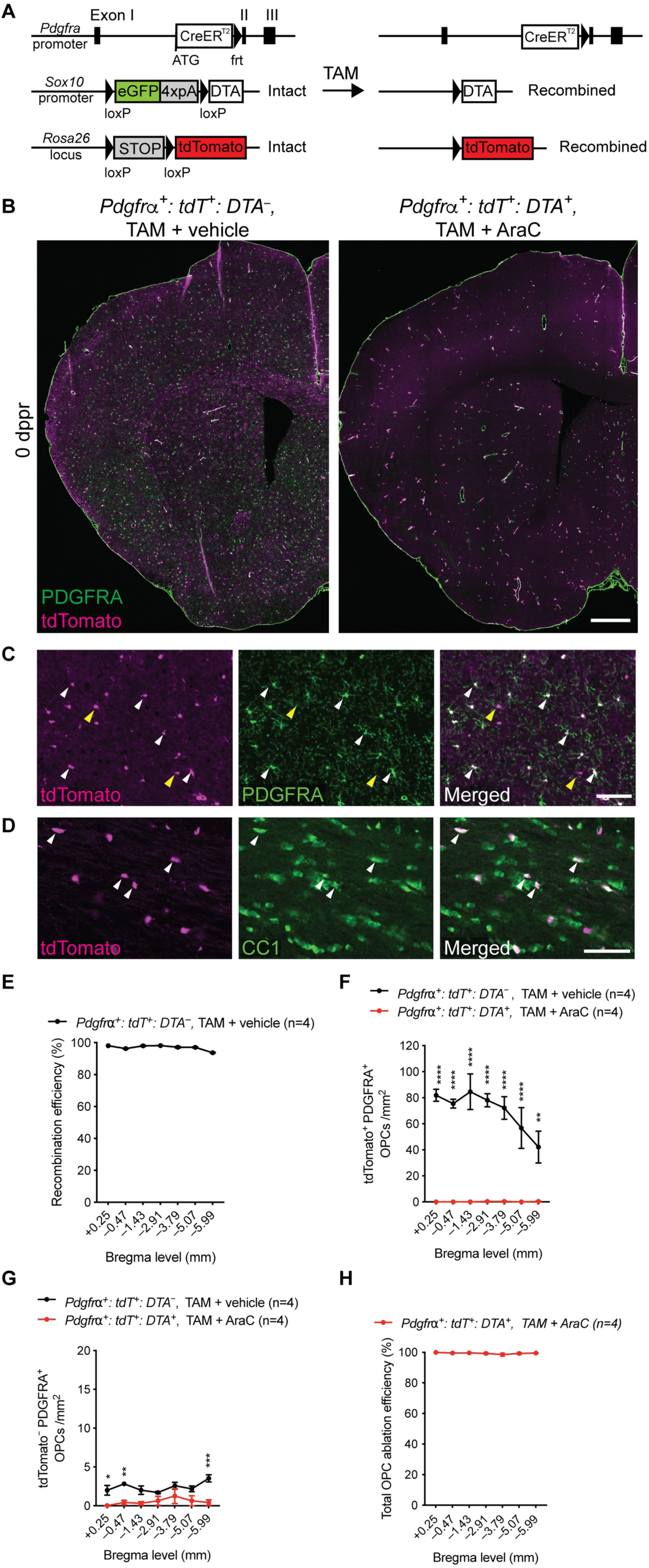
Combined fate-mapping and pharmacogenetic ablation revealed highly efficient and widespread OPC ablation. **A**, Schematic of the transgenic alleles in *Pdgfrα^+^: tdT^+^: DTA^+^* mice. Following TAM administration to *Pdgfrα^+^: tdT^+^: DTA^+^* mice, DTA and tdTomato expression in recombined OPCs was induced after Cre-mediated excision of the *GFP-poly(A)* cassette and the *STOP* cassette, respectively. **B**, Immunohistochemistry against PDGFRA and tdTomato in the rostral forebrain of transgenic mice assessed at 0 dppr. Coronal sections are approximately +0.25 mm anterior of Bregma. Compared to non-ablated controls, OPC-ablated mice exhibited an almost complete OPC loss. **C,D**, tdTomato expression in non-ablated mice (TAM + vehicle administered *Pdgfrα^+^: tdT^+^: DTA^−^* mice) is observed in both PDGFRA^+^ OPCs (**C**, white arrowheads) and in CC1^+^ OLs (**D**, white arrowheads). tdTomato^+^ PDGFRA^−^ cells (**C**, yellow arrowheads) are presumptive fate-mapped OPCs that have differentiated into OLs. **E**, Recombination efficiency of the *Ai14 tdTomato* allele across the brains of TAM + vehicle administered *Pdgfrα^+^: tdT^+^: DTA^−^* mice based on assessment of tdTomato expression among PDGFRA^+^ OPCs. Rostrocaudal position relative to Bregma is indicated on the x-axis. **F**, Cell densities of tdTomato^+^ OPCs in coronal brain sections of non-ablated control versus OPC-ablated mice at 0 dppr. **G**, Densities of tdTomato^−^ OPCs in coronal sections of non-ablated and OPC-ablated mice at 0 dppr. **H**, Percentage depletion of all PDGFRA^+^ OPCs (tdTomato^+^ or tdTomato^−^) in OPC-ablated mice relative to non-ablated controls at 0 dppr. Data represent mean ± SEM. Statistical analysis: two-way ANOVA with Bonferroni’s *post hoc* analysis (**F,G**); \**p*<0.05, \*\**p*<0.01, \*\*\**p*<0.001, \*\*\*\**p*<0.0001. Scale bars, 500 µm (**B**), 100 µm (**C**), 50 µm (**D**).

In non-ablated controls sacrificed at the end of vehicle infusion, 97.0 ± 0.6% of PDGFRA^+^ OPCs were found to express tdTomato (**Figure 2B,C,E**), irrespective of differences in the local density of PDGFRA^+^ OPCs along the rostrocaudal axis of the brain (**Figure 2F**). Cellular morphology was used to discriminate between OPCs and perivascular fibroblast-like cells (VLMCs), the former possessing fine ramified processes whereas the latter were devoid of fine processes and exhibited a circular morphology consistent with vascular localization. We also identified numerous tdTomato^+^ CC1^+^ OLs generated by these fate-mapped OPCs in non-ablated controls (**Figure 2D**). By contrast, virtually no tdTomato^+^ PDGFRA^+^ cells exhibiting typical OPC morphology were detected in the brains of *Pdgfrα^+^: tdT^+^: DTA^+^* mice administered TAM + AraC (**Figure 2B,F, Figure S2E**). Quantitatively, the mean density of tdTomato^+^ OPCs in the AraC-infused brains was 99.7 ± 0.2% lower than that observed in vehicle-infused brains (0.16 ± 0.07 cells/mm^2^ versus 70.1 ± 5.8 cells/mm^2^, *p*<0.0001). We also identified significantly fewer non-recombined (tdTomato^−^) OPCs in ablated mice compared to non-ablated controls (0.53 ± 0.14 cells/mm^2^ versus 2.3 ± 0.2 cells/mm^2^, *p*<0.0001) (**Figure 2G**). When tdTomato^+^ and tdTomato^−^ OPC counts are combined, this equates to a 98.6 ± 0.4% reduction in total OPC density across the entire brain of OPC-ablated mice when compared to non-ablated control mice at 0 dppr (p<0.0001) (**Figure 2H**). By 10 dppr, the mean density of tdTomato^+^ OPCs in ablated mice had increased marginally but remained 97.1 ± 0.6% lower than that observed in vehicle-infused mice (2.00 ± 0.93 cells/mm^2^ versus 70.1 ± 5.8 cells/mm^2^, *p*<0.0001). At this time-point, PDGFRA^+^ cells remained depleted in the cerebrum, but began to repopulate caudoventral regions of the brain, particularly the brain stem (**Figure S3A**). By 20 dppr, PDGFRA^+^ cells were evident in both the brain stem and cerebrum (**Figure S3B**). Collectively, our results demonstrate that the use of genetically encoded DTA targeting cells expressing PDGFRA and SOX10 in conjunction with intracisternal infusion of the anti-mitotic agent AraC results in highly efficient OPC ablation throughout the entire brain that persists for at least 10 dppr after which PDGFRA^+^ cells started to reappear, first in the brain stem then later in the cerebrum.

### Effects of OPC ablation on other cell types in the brain

To assess whether the induction of DTA-mediated apoptosis was restricted to OPCs within the oligodendroglial lineage, we quantified the densities of CC1^+^ mature OLs in coronal brain sections of OPC-ablated and non-ablated controls. Immunohistochemical analyses of brain sections revealed similar OL density between groups (**Figure 3A-C**). As expected, the transient ablation of PDGFRA^+^ NG2^+^ cells resulted in a complete yet temporary disruption in oligodendrogenesis. This was evaluated by quantitating the density of ASPA^+^ EdU^+^ cells in the corpus callosum of OPC-ablated and non-ablated controls that were administered EdU continuously in their drinking water following infusion, until they were perfused at either 11, 20 or 34 dppr (**Figure 3D**). Consistent with this finding, examination of the corpus callosum at 0, 10 and 20 dppr revealed that our pharmacogenetic approach resulted in the loss of both early (PDGFRA^+^ GPR17^−^) and late (PDGFRA^+^ GPR17^+^) OPCs as well as committed oligodendrocyte progenitors (COPs) (PDGFRA^−^ GPR17^+^) that are in the process of transitioning into mature OL for at least 10 dppr before returning to control levels by 20 dppr (**Figure S4A,B**). Despite the transient reduction in oligodendrogenesis, the absolute density of callosal ASPA^+^ oligodendrocytes of OPC-ablated mice was not significantly different from non-ablated controls examined at the same time-point (**Figure 3C**), and no differences in myelin abundance in the corpus callosum were detected (**Figure 3E,F**). Together these findings suggest that OPC ablation resulted in a transient disruption to the production of newborn oligodendroglia but did not affect pre-existing mature myelin-forming OLs.

**Figure 3.**
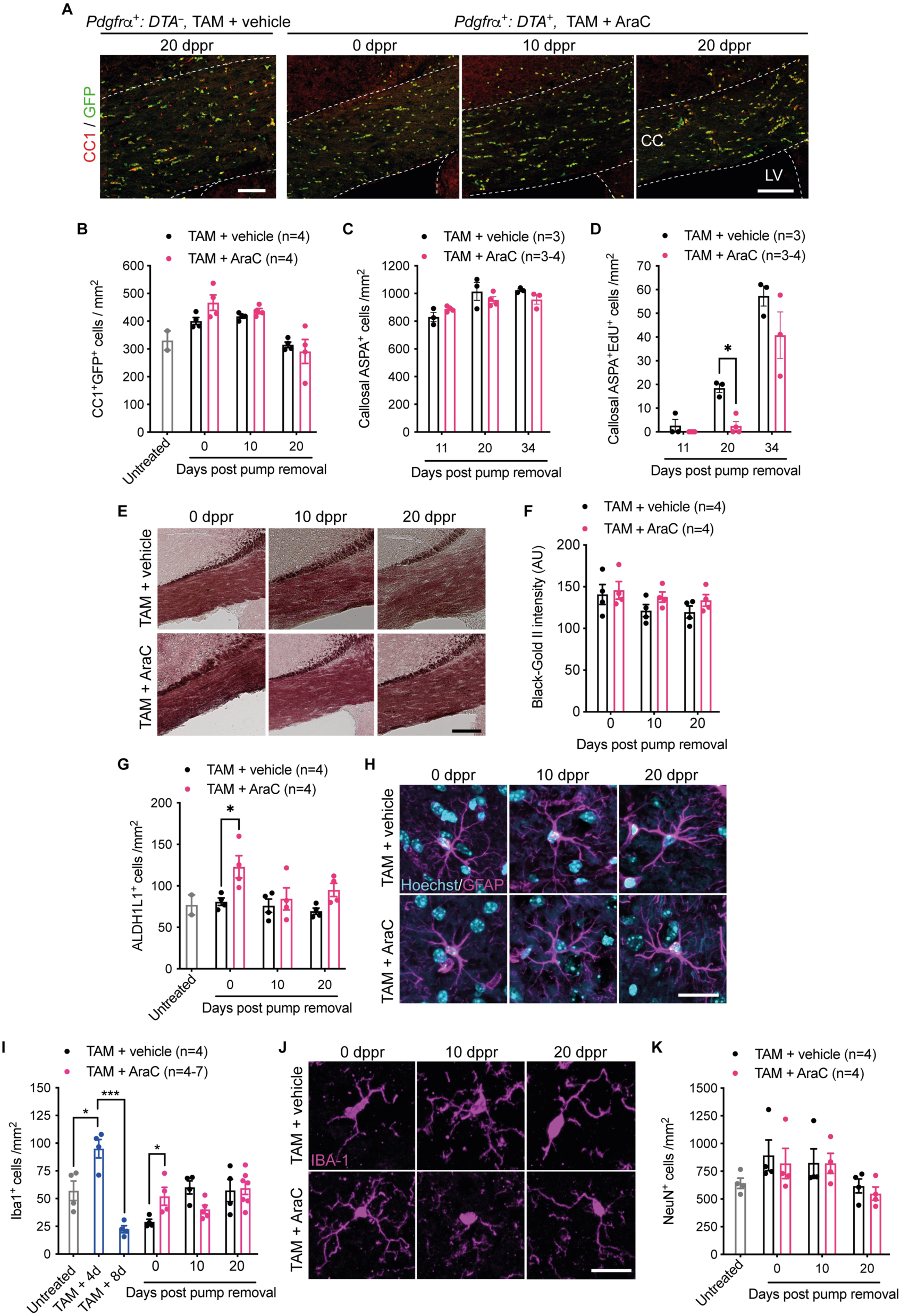
Effect of OPC ablation on other neural cell types. **A**, Coronal sections of the rostral corpus callosum of TAM-administered *Pdgfrα^+^: DTA^+^* mice were immunostained with antibodies against GFP and CC1. **B**, Mean densities of CC1^+^GFP^+^ cells in the rostral segment of the forebrain (gray and white matter combined) of each mouse group. OPC ablation did not change the density of mature OLs at the assessed time-points. **C**, Densities of ASPA^+^ OLs in the corpus callosum remained similar in OPC-ablated versus control mice. **D**, Generation of newborn OLs (ASPA^+^ EdU^+^ cells) after infusion was transiently reduced in OPC-ablated mice. **E**, Black-Gold II-stained coronal sections of the rostral corpus callosum of vehicle- or AraC-infused mice at days 0, 10 and 20 post pump removal. **F**, Quantification of the mean Black-Gold II myelin intensity in the entire corpus callosum revealed no effect of OPC ablation on myelination. **G**, Analysis of ALDH1L1^+^ astrocyte density in the rostral segment of forebrain revealed an overall effect of OPC ablation (p=0.0038) and a significant increase in the number of activated astrocytes was observed at the end of AraC infusion (\**p*=0.0153). **H**, GFAP immunohistochemistry revealed that OPC ablation has minimal effect on astrocyte morphology in the cerebral cortex. **I**, Densities of Iba1^+^ cells in the rostral corpus callosum of *Pdgfrα^+^: DTA^+^* mice that received no treatment (grey bar), TAM alone (blue bars), or either TAM + vehicle (black bars) or TAM + AraC (magenta bars) and were assessed at 0, 10 or 20 dppr (\**p*<0.05, \*\*\**p*<0.001, unpaired two-tailed Student’s *t* test). **J**, Iba1 immunohistochemistry revealed that OPC ablation transiently reduced microglial process complexity in the cerebral cortex. **K**, Densities of NeuN^+^ cells in the rostral cerebral cortex revealed no effect of OPC ablation on cortical neurons. Sample size, *n=*3-7 mice per group, mean ± SEM (**B**-**D**,**F**,**G**,**I**,**K**). CC, corpus callosum; LV, lateral ventricle. Scale bars, 200 µm (**A**), 150 µm (**E**), 25 µm (**H**) and 20 µm (**J**).

Next, we turned our examination to other glial cell types in the CNS of OPC-ablated mice. The density of ALDH1L1^+^ astrocytes was elevated at 0 dppr in *Pdgfrα^+^: DTA^+^* mice administered TAM + AraC and returned to control levels by 10 dppr (**Figure 3G**). The transient increase in astrocyte density was not accompanied by any notable change in the expression of the intermediate filament protein GFAP, a marker of astrocyte activation (**Figure 3H**). Similarly, the morphology of GFAP^+^ astrocytes was equivalent in OPC-ablated and non-ablated controls, although soma size and the number and length of processes increased marginally in both groups with time post-infusion (**Figure S5B-E**). In terms of microglial response, we observed a transient increase in the density of Iba1^+^ microglia in *Pdgfrα^+^: DTA^+^* mice 4 days after final TAM administration and at the end of AraC infusion (0 dppr) when compared to non-ablated controls (0 dppr) which normalized by 20 dppr (**Figure 3I**). Morphological analysis of Iba1^+^ microglia revealed that OPC-ablated mice exhibited changes in process complexity and somal area over time. At 0 dppr, microglia exhibited an increase in the number of secondary processes, reflective of a more active (hyper-ramified) state. Conversely, by 10 dppr, there was a significant reduction in both primary and secondary processes, as well as process length, suggesting the presence of ameboid or dystrophic microglia (**Figure 3J, Figure S6A-D**). These morphological changes were not associated with any significant shifts in the percentage of microglia that expressed the M1- or M2-associated markers CD16/CD32 or CD206, respectively (**Figure S6E,F**), although we noted that the level of expression of CD16/CD32 was elevated at 0 dppr in OPC-ablated mice before subsequently declining. Together, these data suggest that OPC ablation induced a modest and transient neuroinflammatory response that had largely resolved by 20 dppr.

To assess whether extensive OPC ablation led to neuronal cell death through neuroinflammation as demonstrated in a previous study (Nakano *et al*., 2017), NeuN^+^ neurons were quantified in the cerebral cortex where PDGFRA^+^ cells remained depleted until 20 dppr. Interestingly, the densities of cortical neurons in the cerebral cortex were comparable between groups at all time-points (**Figure 3K**), irrespective of cortical layer (data not shown), suggesting that OPC ablation does not compromise the viability of cortical neurons.

### PDGFRA^+^ cells started to repopulate the cerebrum from 12 dppr but did not derive from the OPC lineage

To examine the kinetics and anatomical origin of PDGFRA^+^ cells that repopulated the cerebrum following OPC ablation, additional cohorts of TAM + AraC administered *Pdgfrα^+^: tdT^+^: DTA^+^* mice were sacrificed at 0, 12, 18, 20 and 34 dppr. TAM + vehicle administered *Pdgfrα^+^: tdT^+^: DTA^−^* mice served as non-ablated controls (**Figure 4A**). In the corpus callosum of OPC-ablated mice, we confirmed highly efficient ablation of tdTomato^+^ PDGFRA^+^ OPCs at 0 dppr (**Figure 4B**). However, by 20 dppr, we observed high densities of PDGFRA^+^ cells that did not express tdTomato, indicating that the vast majority of these cells do not derive from surviving tdTomato^+^ OPCs (**Figure 4B, Figure S7A-C**).

**Figure 4.**
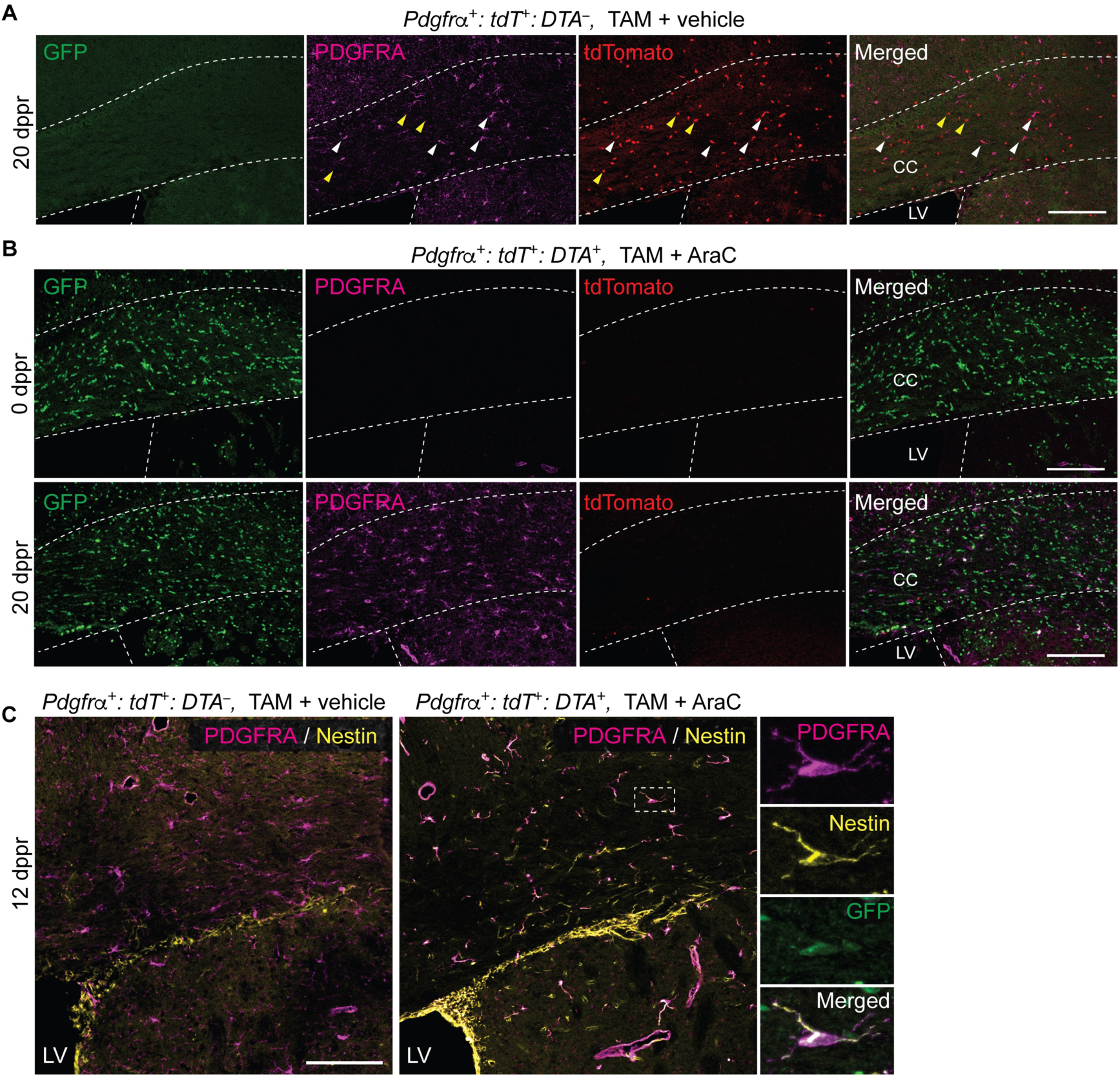
PDGFRA^+^ cells that repopulate the corpus callosum following OPC ablation are not derived from non-ablated fate-mapped OPCs. **A,B**, Immunohistochemical detection of GFP^+^, PDGFRA^+^ and tdTomato^+^ cells in coronal sections of the anterior forebrain of non-ablated versus OPC-ablated mice at 20 dppr. In vehicle-infused controls that do not carry the *Sox10-DTA* allele, the vast majority of PDGFRA^+^ OPCs co-express tdTomato (white arrowheads), reflecting high recombination efficiency of the *Ai14 tdTomato* allele (**A**). In addition, the tdTomato^+^ PDGFRA*^−^* cells (yellow arrowheads) are presumptive fate-mapped OPCs that have differentiated into OLs following TAM administration. **B,** Fate-mapped OPCs expressing PDGFRA and tdTomato were not detected at 0 dppr in TAM + AraC administered *Pdgfrα^+^: tdT^+^: DTA^+^* mice. By 20 dppr, PDGFRA^+^ cells predominately repopulated the corpus callosum proximal to the SVZ and did not express tdTomato suggesting that they do not derive from the OPC lineage. **C**, Virtually all PDGFRA^+^ cells in the corpus callosum proximal to the V-SVZ expressed the NPC marker Nestin at the initial stage of regeneration at 12 dppr of AraC. High magnification of the boxed region reveals a GFP^+^ PDGFRA^+^ cell expressing Nestin. CC, corpus callosum; LV, lateral ventricle. Data represent mean ± SEM. CC, corpus callosum; LV, lateral ventricle. Scale bars, 40 µm (**A,B**) and 100 µm (**C**).

PDGFRA^+^ cells repopulating the cerebrum first appeared at 12 dppr in a region of the corpus callosum adjacent to the V-SVZ. Unlike PDGFRA^+^ cells found in the corpus callosum of non-ablated controls, those in OPC-depleted mice co-expressed Nestin, an intermediate filament protein normally expressed by cells in the V-SVZ (**Figure 4C**). Newly generated PDGFRA^+^ NG2^+^ cells in OPC-depleted mice also expressed GFP, indicating that the *Sox10-DTA* allele in these cells was in a non-recombined state, indicating that they do not derive from recombined OPCs (**Figure S4C**). In addition, repopulating PDGFRA^+^ cells were EdU^+^, indicating that they were born after AraC infusion (data not shown) and many possessed a unipolar or bipolar morphology consistent with migratory activity (Tsai et al., 2009).

The density of PDGFRA^+^ cells in the rostral cerebrum of OPC-depleted mice returned to levels similar to that of non-ablated controls in a spatiotemporally defined manner, normalizing first in the region of the corpus callosum adjacent to the V-SVZ at 12 dppr (**Figure 5B**), followed by the midline corpus callosum at 18 dppr (**Figure 5C**), and later in the cerebral cortex at 20 dppr (**Figure 5D**). Other regions of the rostral cerebrum of OPC-depleted mice exhibited different latencies for PDGFRA^+^ cell density to return to control levels (**Figure 5E-H**). Overall, the mean density of PDGFRA^+^ cells in the rostral cerebrum returned to levels similar to that of non-ablated controls by 20 dppr whereas caudal regions of the cerebrum remained deficient at the same time-point (**Figure S3C**).

**Figure 5.**
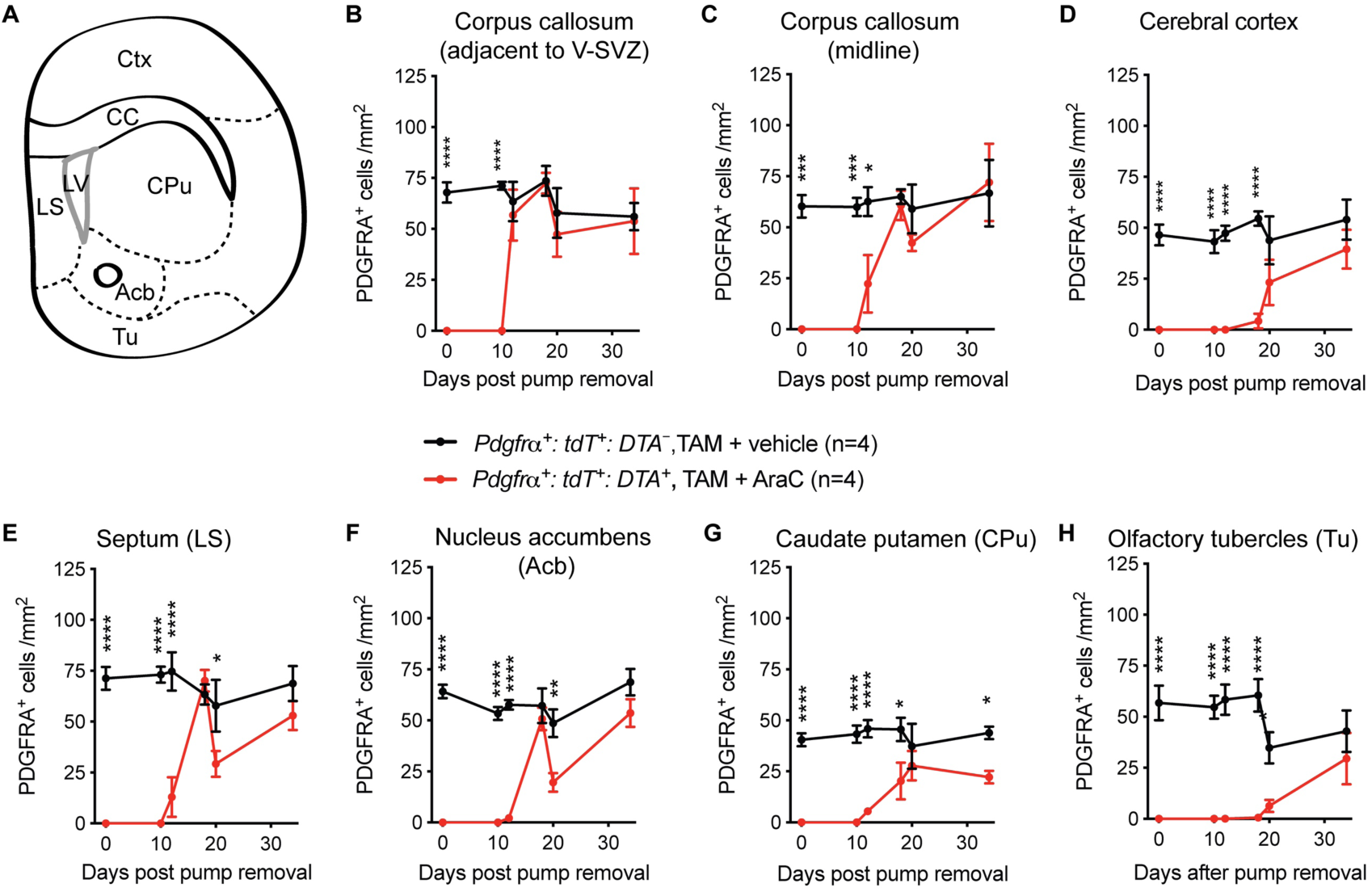
Spatiotemporal profile of repopulation of newly-generated PDGFRA^+^ cells in the anterior forebrain of OPC-depleted mice. **A,** Schematic diagram depicting various brain regions within which PDGFRA^+^ cell densities were quantitated. **B-H**, Densities of PDGFRA^+^ cells in the region of the corpus callosum adjacent to the V-SVZ, at the midline of the corpus callosum, in the cerebral cortex, septum, nucleus accumbens, caudate putamen and olfactory tubercles from 0-34 dppr revealed distinct spatiotemporal patterns of regeneration of PDGFRA^+^ cells (n=4 mice per group, mean ± SEM; *p<0.05, **p<0.01, \*\*\**p*<0.001, \*\*\*\**p*<0.0001, two-way ANOVA with Bonferroni’s *post hoc* analysis).

Collectively these data demonstrate that ablation of 98.6 ± 0.4% of OPCs was followed by a late onset regenerative response resulting in the repopulation of PDGFRA^+^ cells. The finding that the vast majority of repopulating PDGFRA^+^ cells in the cerebrum were tdTomato^−^ and GFP^+^ is inconsistent with the notion that PDGFRA^+^ cells arise through proliferative expansion of surviving OPCs. Rather, the spatiotemporal pattern of PDGFRA^+^ cell regeneration, emerging first in a region of the corpus callosum adjacent to the V-SVZ, whilst co-expressing the V-SVZ marker Nestin, raised the possibility that NPCs within the V-SVZ could be the primary source of PDGFRA^+^ cells that repopulated this region of the brain following OPC ablation.

### Regeneration of V-SVZ-derived NPCs after AraC infusion

To investigate whether NPCs in the V-SVZ could serve as a reservoir to regenerate PDGFRA^+^ cells, we examined the response of NPCs to pharmacogenetic ablation of OPCs. A 6-day infusion of 2% AraC onto the surface of the brain was previously demonstrated to eliminate rapidly dividing cells in the V-SVZ (Doetsch et al., 1999a). The subsequent activation and proliferation of quiescent neural stem cells in the V-SVZ resulted in complete regeneration of the neurogenic niche including transit-amplifying cells and neuroblasts within 10 days after AraC withdrawal (Doetsch et al., 1999b). To assess the regeneration of NPCs in TAM + AraC administered *Pdgfrα^+^: DTA^+^* mouse brains, mice were given EdU via their drinking water beginning immediately after pump removal and continuing until they were perfused 10 days later (**Figure 1H**). Compared to control mice, DCX^+^ neuroblasts were virtually absent from the V-SVZ of AraC-infused mice at 0 dppr (**Figure 6A**). At 10 dppr, the number of DCX^+^ cells in the V-SVZ of AraC-treated animals was similar to that of vehicle-treated controls (**Figure 6A,B**). In addition, these DCX^+^ cells were positive for EdU (**Figure 6C**), indicating that they were born after AraC infusion. We conclude that the AraC infusion ablated rapidly dividing NPCs in the V-SVZ but these cells were completely regenerated by 10 dppr.

**Figure 6.**
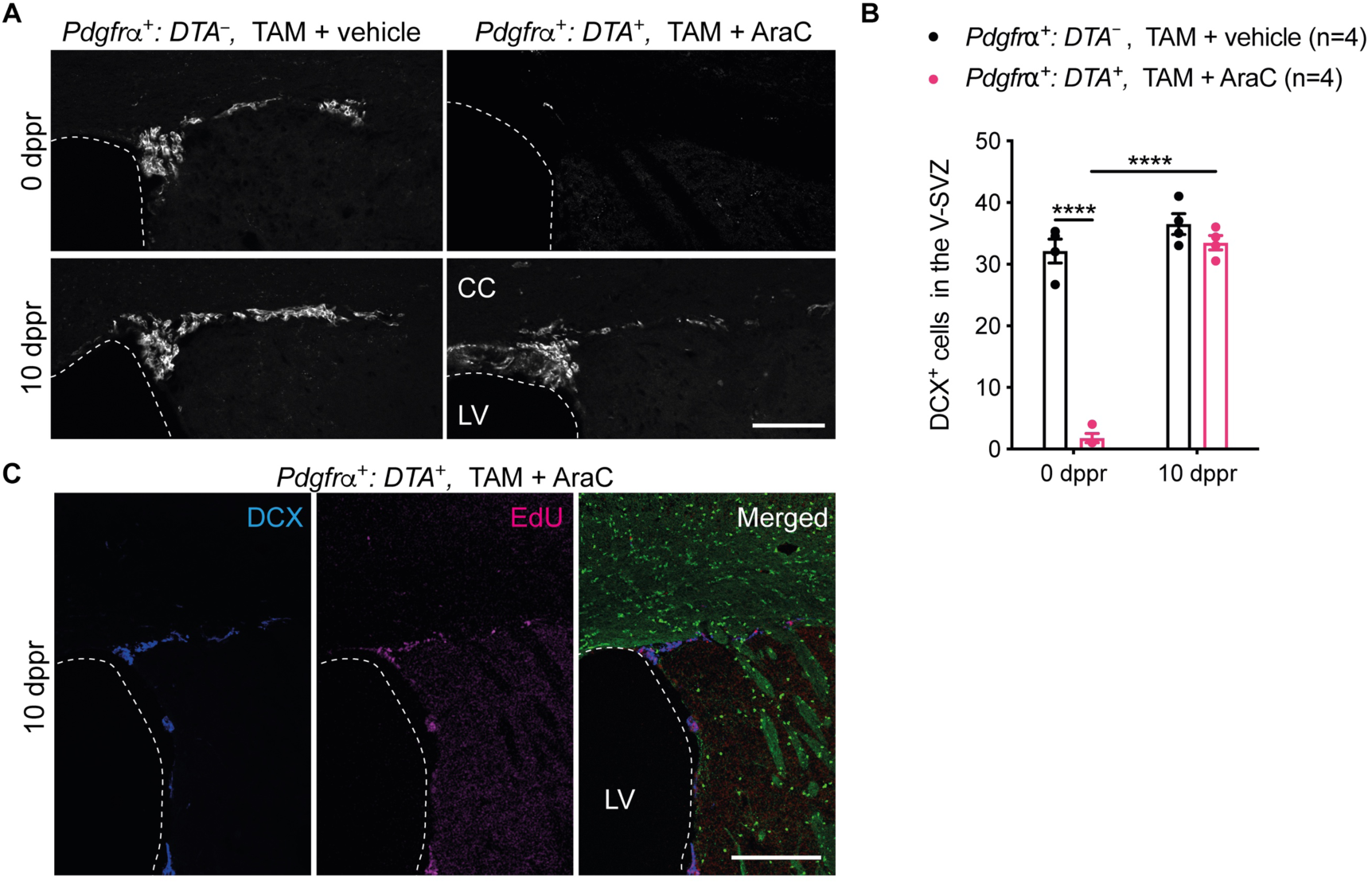
AraC infusion resulted in the transient depletion of neuroblasts in the V-SVZ followed by their complete regeneration within 10 dppr. **A**, Expression of the neuroblast marker doublecortin (DCX) in the V-SVZ of *Pdgfrα^+^: DTA^+^* mice administered either TAM + vehicle or TAM + AraC and assessed at 0 or 10 dppr. **B**, Densities of DCX^+^ cells in the V-SVZ of TAM + vehicle or TAM + AraC administered *Pdgfrα^+^: DTA^+^* mice assessed at 0 or 10 dppr (*n=*4 mice per group, mean ± SEM). (\*\*\*\**p*<0.0001, two-way ANOVA with Bonferroni’s *post hoc* analysis.) **C**, Coronal sections of the V-SVZ immunolabeled with antibodies against GFP and DCX, and stained for EdU, which was administered to transgenic mice for 10 dppr. EdU^+^ neuroblasts defined by co-expression of DCX were restricted to the V-SVZ. CC, corpus callosum; LV, lateral ventricle. Scale bars, 100 μm (**A**) and 200 μm (**C**).

### V-SVZ-derived NPCs contributed to the regeneration of PDGFRA^+^ cells in the dorsal anterior forebrain after OPC ablation

Our earlier examination of OPC-ablated mice at 12 dppr had revealed that callosal PDGFRA^+^ cells adjacent to the V-SVZ co-expressed Nestin, a well-established marker of NPCs in the V-SVZ, whereas callosal OPCs in the vehicle-infused controls did not (**Figure 4C**). We posited that repopulating PDGFRA^+^ cells could derive from progenitors located within the V-SVZ. In support of this possibility, we found that in the corpus callosum of OPC-ablated mice examined at 20 dppr, PDGFRA^+^ NG2^+^ cells co-expressed GFP indicating that the *Sox10-GFP* allele was in a non-recombined state (**Figure S4C**). We also found that PDGFRA^+^ NG2^+^ cells in the dorsolateral corner as well as in the dorsal and lateral walls of the V-SVZ co-labelled with EdU which was provided to the mice following AraC infusion (data not shown).

To confirm that NPCs generate PDGFRA^+^ cells that migrate into the cerebrum following OPC ablation, we developed a novel Dre recombinase-based viral approach for the genetic fate mapping of V-SVZ-derived NPCs (**Figure S8A-D**). Two weeks prior to pharmacogenetic ablation of OPCs, we injected *Pdgfrα^+^: tdT^+^: DTA^+^* mice with high-titer lentiviruses to transduce NPCs in the V-SVZ (**Figure 7A**). This resulted in the expression of a Myc-tagged membrane-targeted mKate2 fluorescent reporter protein by a subset of cells in the V-SVZ (**Figure S8E-I**). At 20 dppr, we detected the expression of mKate2 in a subpopulation of newly-generated PDGFRA^+^ cells in the corpus callosum, indicating that they had derived from the NPC lineage (**Figure 7B,C**). The V-SVZ origin of newly-generated PDGFRA^+^ cells was corroborated by simultaneous ablation of both parenchymal OPCs and oligodendrogenic NPCs using *Nestin-CreER^T2+/–^; Pdgfrα-CreER^T2+/–^; Sox10-DTA^+/−^*transgenic mice (denoted hereafter as *Nestin^+^: Pdgfra^+^: DTA^+^* mice) (**Figure 7D**). The persistent absence of PDGFRA^+^ cells in the cerebrum of parenchymal OPC and oligodendrogenic NPC co-ablated mice at 20 dppr after AraC withdrawal indicates that regenerating PDGFRA^+^ cells in the dorsal cerebrum of OPC-ablated mice derive from Nestin-expressing NPCs residing in the V-SVZ (**Figure 7E,F, Figure S9A**). Of note, *Nestin^+^: Pdgfra^+^: DTA^+^*mice exhibited hydrocephalus, which we believe reflects a degree of TAM-independent recombination in SOX10^+^ Nestin^+^ neural crest cells during development (see *Supplementary Information*). Despite evidence of hydrocephalus, we observed a similar density of neurogenic NPCs expressing DCX in the lateral walls and dorsolateral corner of the V-SVZ of TAM + AraC administered *Nestin^+^: Pdgfra^+^: DTA^+^* mice when compared to non-ablated controls (**Figure S9C,D**). Thus, while neuroblast production persists in TAM + AraC administered *Nestin^+^: Pdgfra^+^: DTA^+^* mice, PDGFRA^+^ cells are not viable. Collectively, these data indicate that PDGFRA^+^ cells that repopulate the dorsal cerebrum of OPC-depleted mice beginning 12 dppr arise from oligodendrogenic NPCs residing in the V-SVZ.

**Figure 7.**
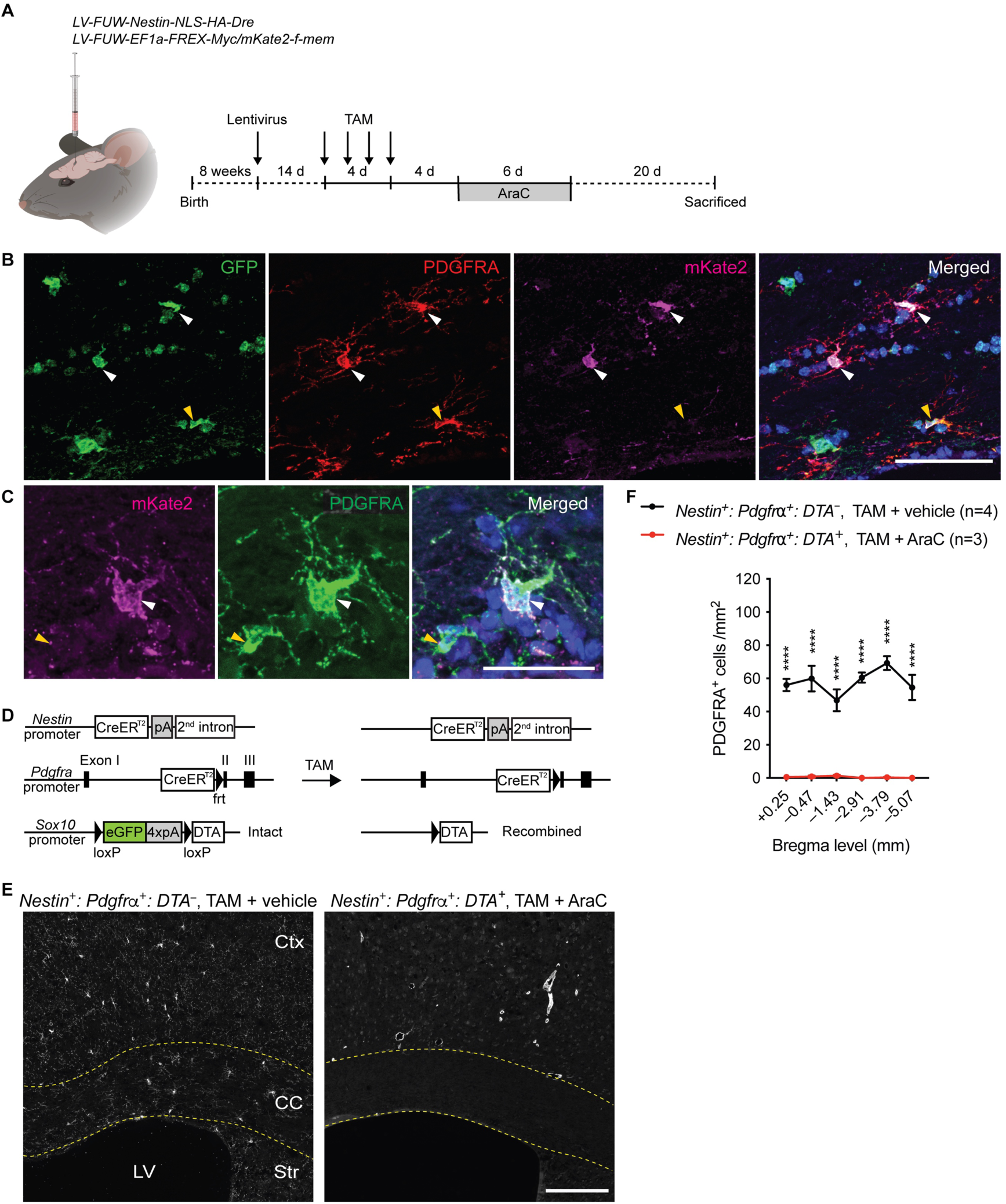
V-SVZ-derived NPCs were responsible for the regeneration of new PDGFRA^+^ cells in the anterior/dorsal forebrain after OPC ablation. **A**, Schematic diagram and timeline of intraventricular injection of lentiviruses to map the fate of V-SVZ-derived NPCs in adult *Pdgfrα^+^: tdT^+^: DTA^+^* mice. Schematic diagram was created using BioRender. **B,C**, In *Pdgfrα^+^: tdT^+^: DTA^+^,* TAM + AraC mice assessed at 20 dppr, immunostaining with antibodies against PDGFRA, GFP and mKate2 revealed some transduced newly-generated PDGFRA^+^ cells (white arrowheads) expressing membrane-targeted mKate2 in the corpus callosum proximal to the V-SVZ, suggesting they were derived from the NPC lineage. Non-transduced PDGFRA^+^ cells are indicated with yellow arrowheads. **D,** Schematic of transgenic alleles in *Nestin^+^: Pdgfrα^+^: DTA^+^* mice. Following TAM administration, DTA expression in recombined OPCs and oligodendrogenic NPCs is induced after Cre-mediated excision of the *GFP-poly(A)* cassette contained within the *Sox10-DTA* allele. **E,** Immunohistochemical detection of PDGFRA in the rostral cerebrum of non-ablated controls (*Nestin^+^: Pdgfrα^+^: DTA^−^,* TAM + vehicle mice) and in parenchymal OPC and oligodendrogenic NPC co-ablated mice (*Nestin^+^: Pdgfrα^+^: DTA^+^,* TAM + AraC) at 20 dppr. **F,** Densities of PDGFRA^+^ cells in the cerebrum of non-ablated controls and in parenchymal OPC and oligodendrogenic NPC co-ablated mice at 20 dppr. (*n=*3-4 mice per group, mean ± SEM). (\*\*\*\**p*<0.0001, two-way ANOVA with Bonferroni’s *post hoc* analysis.) Ctx, cortex; CC, corpus callosum; LV, lateral ventricle; Str, striatum. Scale bars, 60 µm (**B**), 30 µm (**C**) and 100 µm (**E**).

## Discussion

The development of methods to conditionally ablate parenchymal OPCs in the adult mouse CNS provides a powerful experimental paradigm to explore their function. While approaches to ablate adult OPCs have been described (Birey *et al*., 2015; Dang *et al*., 2019; Hughes *et al*., 2013; Irvine and Blakemore, 2007; Nakano *et al*., 2017; Robins *et al*., 2013), the techniques established to date have not allowed for the complete depletion of OPCs throughout the brain. In the absence of complete OPC ablation, surviving OPCs enter the cell cycle and rapidly regenerate their population as a homeostatic response, complicating the interpretation of results. To avert rapid regeneration by surviving OPCs, we have developed a novel pharmacogenetic approach that allows for the highly efficient ablation of OPCs throughout the brain. This approach consists of genetic elements permitting TAM-dependent induction of DTA expression exclusively in OPCs followed by a pharmacological intervention to prevent repopulation by OPCs that avert genetic targeting. Genetic control of OPC ablation is achieved by crossing the *Pdgfra-CreER^T2^* and *Sox10-lox-GFP-STOP-lox-DTA* transgenic mouse lines. Double transgenic mice enable highly specific ablation of OPCs at a prescribed time-point due to the dependence of DTA expression upon three levels of regulation: 1) activity of the *Pdgfra* promoter, 2) activity of the oligodendroglial-specific *Sox10* promoter, and 3) temporal control provided by TAM-dependent regulation of CreER^T2^ activity. However, delivery of TAM alone to *Pdgfrα^+^: DTA^+^* mice was insufficient for complete or durable OPC ablation, due to the rapid proliferation of residual surviving OPCs. Since cumulative toxicity precludes delivery of TAM beyond 4 days, following TAM delivery, mice were infused intracisternally with the anti-mitotic drug AraC to prevent OPC that escape genetic targeting from re-entering the cell cycle and repopulating the brain. In isolation, intracisternal infusion of AraC to wild-type mice did not reduce OPC density, consistent with the fact that only a fraction of OPCs in the healthy adult CNS are dividing at a given time and the homeostatic control mechanisms that maintain OPC density when OPCs are sparsely depleted (Hughes *et al*., 2013). However, the combined use of both genetic and pharmacological approaches eliminated 98.6 ± 0.4% of all OPCs throughout the brain without causing any overt adverse effects on the health of animals.

The combined pharmacogenetic approach that we have developed to target OPCs is anticipated to result in the death of cells expressing both PDGFRA and SOX10 and mitotic cells exposed to AraC. By examining the densities of various non-OPC cell types in the brains of OPC ablated mice, our data suggest that while OPCs, as well as committed oligodendrocyte progenitors (COPs) (PDGFRA^−^ GPR17^+^) and DCX^+^ cells in the V-SVZ are depleted, the density of mature oligodendrocytes, astrocytes and neurons remains unchanged. Significantly, however, the ablation of OPCs is anticipated to trigger a broad range of direct and indirect effects upon other neural cell types that are not necessarily reflected by a change in cell density. For instance, the activation of microglia to facilitate phagocytosis of cellular debris of OPCs undergoing cell death is expected (Marquez-Ropero et al., 2020). Consistent with this possibility, we identified an initial increase and subsequent decrease in the density of microglia over the course of OPC ablation and subsequent regeneration. We also observed changes in microglial phenotype suggesting that OPC ablation alters the activation status of innate immune cells in the CNS. Given these observations, the effect that changes in microglial density and activation status have upon behavioral, cellular and molecular readouts should be taken into consideration in the future application of the model. One approach to address the role that microglia exert in this model would be to deplete microglia via inhibition of the colony-stimulating factor 1 receptor (CSF1R) (Green et al., 2020), in the context of OPC ablation, to discern the immune-mediated effects of OPC ablation from primary effects caused by depletion of OPCs.

Despite the highly efficient ablation of OPCs throughout the brain, PDGFRA^+^ NG2^+^ cells were eventually regenerated, beginning from around 12 dppr. In this context, it is important to note that AraC infusion also ablated almost all DCX^+^ neuroblasts in the V-SVZ before their return to normal density by 10 dppr. The transient depletion of DCX^+^ cells in the V-SVZ is consistent with previous reports demonstrating that AraC-mediated elimination of rapidly dividing transit-amplifying cells and neuroblasts in the V-SVZ activates quiescent neural stem cells to re-enter the cell cycle, leading to complete regeneration of neuroblasts in the V-SVZ within 10 days following AraC withdrawal (Doetsch *et al*., 1999a; Doetsch *et al*., 1999b). The kinetics of neuroblast regeneration in the V-SVZ aligned closely with the spatiotemporal pattern of PDGFRA^+^ NG2^+^ cell regeneration in the cerebrum. First, we noted that recombined (tdTomato^+^) OPCs did not contribute to the regeneration of PDGFRA^+^ cells. Rather, we showed that non-recombined (GFP^+^) PDGFRA^+^ cells arose first in the vicinity of the V-SVZ and expressed Nestin, a known V-SVZ marker shortly after DCX^+^ cell density returned to control levels after ablation. This observation could be explained by two distinct possibilities. Our *a priori* view was that this reflected recruitment of oligodendrogenic NPCs from the V-SVZ that maintain their expression of Nestin for some time after they migrate out of the V-SVZ, given the known capacity of the V-SVZ to generate PDGFRA^+^ NG2^+^ cells in health and disease (Menn et al., 2006; Xing et al., 2014). An alternate possibility is that Nestin^+^ GFP^+^ PDGFRA^+^ cells observed at 12 dppr reflect non-recombined OPCs that survive pharmacogenetic ablation and transiently upregulate Nestin in response to neuroinflammatory cues, as has been described for astrocytes under ischemic conditions (Duggal et al., 1997; Lin et al., 1995). Countering this alternate possibility, we found that lentiviral-mediated labeling of Nestin-expressing cells in the V-SVZ prior to OPC ablation allowed us to identify mKate2^+^ PDGFRA^+^ cells in the corpus callosum following OPC ablation, providing additional evidence to support the V-SVZ origin of repopulating PDGFRA^+^ cells in the cerebrum. Whilst this experiment demonstrates that NPCs lining the lateral ventricles can give rise to PDGFRA^+^ cells during the regenerative phase that follows OPC ablation, the limited efficiency of this viral labeling approach cannot account for all PDGFRA^+^ cells that are regenerated in the cerebrum. We addressed this issue by adopting a co-ablation strategy that directs Sox10-dependent DTA expression to both the NPC and OPC lineages. Using this approach, we found that the co-ablation of both parenchymal OPCs and oligodendrogenic NPCs resulted in failure to regenerate PDGFRA^+^ cells in the dorsal cerebrum, providing robust evidence that repopulating PDGFRA^+^ cells in the cerebrum derive from the NPC lineage.

A potential caveat of the co-ablation strategy that warrants consideration is the fidelity with which the *Nestin-CreER^T2^* driver is restricted to NPCs. Let us first consider the possibility that the PDGFRA^+^ cells that we observe during early repopulation reflect surviving OPCs that express Nestin ectopically under inflammatory conditions rather than cells of NPC origin. If this were the case, ablation of Nestin-expressing OPCs that survive pharmacogenetic ablation in *Nestin^+^: Pdgfra^+^: DTA^+^* mice would require 100% of these putative Nestin-expressing OPCs to exhibit TAM-independent recombination of the *Sox10-DTA* allele. We were able to exclude this possibility by demonstrating that TAM-independent recombination in NPCs with a history of up to 10 weeks of postnatal Nestin expression occurs with a frequency of less than 1% (**Figure S9G**). In light of these findings, the most parsimonious explanation is that following TAM gavage in *Nestin^+^: Pdgfrα^+^: DTA^+^* mice, the recombined *Sox10-DTA* allele starts to be expressed by NPCs upon their specification to an oligodendroglial fate, which induces DTA-mediated apoptosis of oligodendrogenic NPCs.

Our findings expand on previous work demonstrating that NPCs originating in the V-SVZ give rise to oligodendrogenic progenitors, both under normal physiological conditions and in response to demyelination of proximal white matter tracts (Brousse et al., 2015; Menn *et al*., 2006; Samanta et al., 2015; Xing *et al*., 2014). Another study, in which mice undergo global *Pdgfra* inactivation resulting in OPC depletion, suggested that repopulation of OPCs occurred by expansion of OPCs originating from immature Nestin-expressing cells activated in the meninges and brain parenchyma and from OPCs that escape *Pdgfra* inactivation (Dang *et al*., 2019). However, spatiotemporal examination of PDGFRA^+^ NG2^+^ cell regeneration after OPC ablation in our model does not provide evidence to support a meningeal origin of newly-generated PDGFRA^+^ NG2^+^ cells (**Figure S9E,F**).

Our finding that oligodendrogenic progenitors residing in the V-SVZ play a significant role in the regeneration of PDGFRA^+^ NG2^+^ cells in the anterior dorsal forebrain following OPC ablation warrants further investigation in future studies. First, further refinement of the model to eliminate the contribution of V-SVZ cells to this homeostatic response could provide an even greater window of opportunity to explore the long-term consequences of OPC ablation upon brain function. Second, the model provides the means to characterize the similarities and differences between OPCs and V-SVZ-derived oligodendrogenic progenitors regarding their response to neural injury and their contribution to myelin regeneration and adaptive myelination. Third, a comprehensive analysis of the molecular mechanisms that trigger the V-SVZ to produce vast numbers of oligodendrogenic progenitors in the context of OPC ablation could provide clues to pharmacological approaches to potentiate the contribution of V-SVZ-derived oligodendrogenic progenitors to remyelination.

In the caudoventral brain, we identified extensive repopulation of PDGFRA^+^ cells following ablation, which preceded that observed in the cerebrum. Whilst we observed a very high recombination efficiency of the lox-STOP-lox-tdTomato allele in *Pdgfrα^+^: tdT^+^: DTA^−^* mice, namely 97%, we cannot exclude the possibility that a very small number of OPCs that escape recombination of the *Sox10-lox-STOP-GFP-lox-DTA* allele may also escape recombination of the *lox-STOP-lox-tdTomato* (Ai14) allele and also survive AraC infusion. This may be the case in the brain stem where the majority of non-ablated OPCs were identified. Notably, in the brainstem, repopulation of PDGFRA^+^ cells is associated with the appearance of PDGFRA^+^ cells that express tdTomato and GFP indicating that they likely originate by clonal expansion of pre-existing OPCs in which the *lox-STOP-lox-tdTomato* but not the *Sox10-lox-STOP-GFP-lox-DTA* allele had recombined. This is consistent with the known high recombination efficiency of the *lox-STOP-lox-tdTomato* (Ai14) allele.

Although we have not explored the origin of newly-generated PDGFRA^+^ cells that emerge in the brain stem, we note that both non-recombined tdTomato*^−^* PDGFRA^+^ cells and recombined tdTomato^+^ PDGFRA^+^ cells contribute to OPC regeneration in the brain stem (**Figure S3D**). These data suggest that some OPCs in the brain stem survive the OPC ablation protocol. This also appears to be the case in the optic nerve and spinal cord, which exhibit robust OPC depletion (**Figure S7D-G**). In the brain stem, it is plausible that repopulating tdTomato*^−^* PDGFRA^+^ cells could arise from immature progenitors residing within alternate stem cell niches located within the brain stem. An oligodendrogenic niche has recently been described in the median eminence of the hypothalamus bordering the third ventricle (Zilkha-Falb et al., 2020). Other putative stem cell niches include the ependymal and subependymal zones of the third and fourth ventricles (Lin and Iacovitti, 2015). Notably, we identified high densities of PDGFRA^+^ cells around the aqueduct (**Figure S3B**) and the fourth ventricle (data not shown) at 10 dppr. It thus seems likely that the homeostatic regeneration of PDGFRA^+^ NG2^+^ cells following OPC depletion is mediated by distinct populations of progenitors in the cerebrum and brain stem.

Collectively, our mouse model of conditional OPC ablation provides a long-sought-after methodology to eliminate OPCs in the adult mouse CNS. The model now provides the opportunity to explore the role that OPCs play in CNS homeostasis by probing the consequences of their ablation through various approaches including but not limited to single-cell RNA sequencing and proteomic analysis to identify the transcriptional and post-translational changes of OPC ablated and non-ablated mice.

## Methods

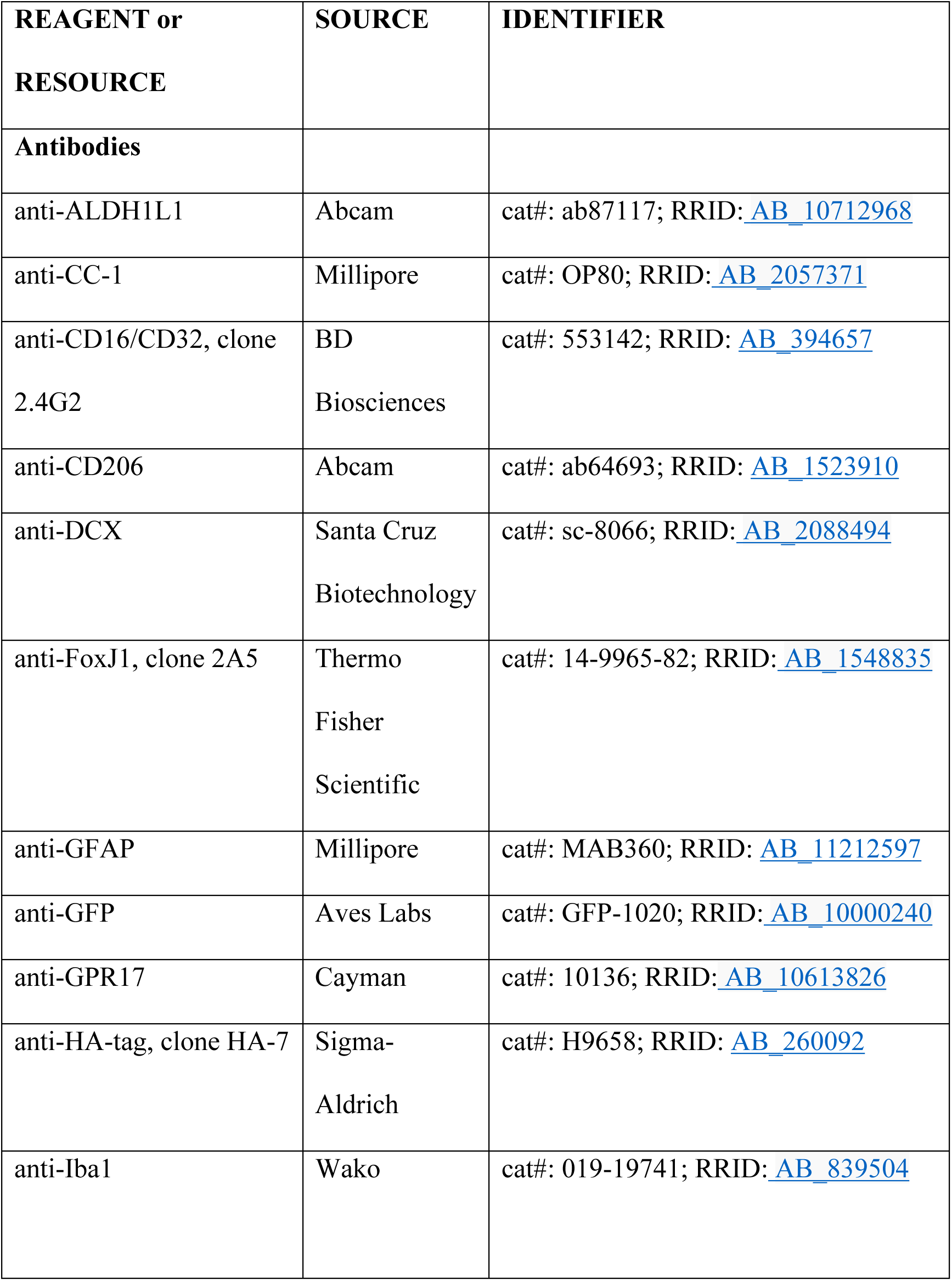
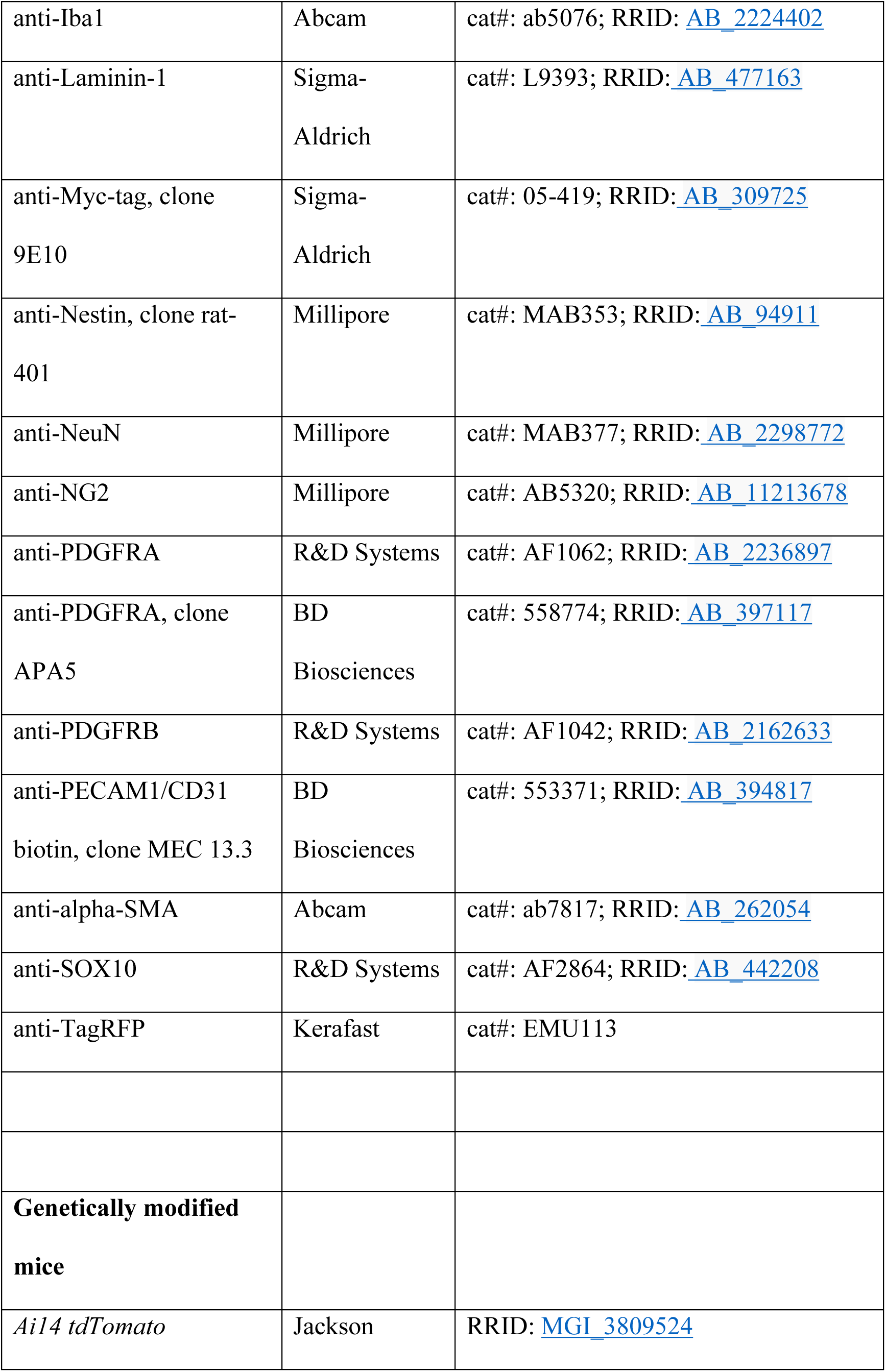
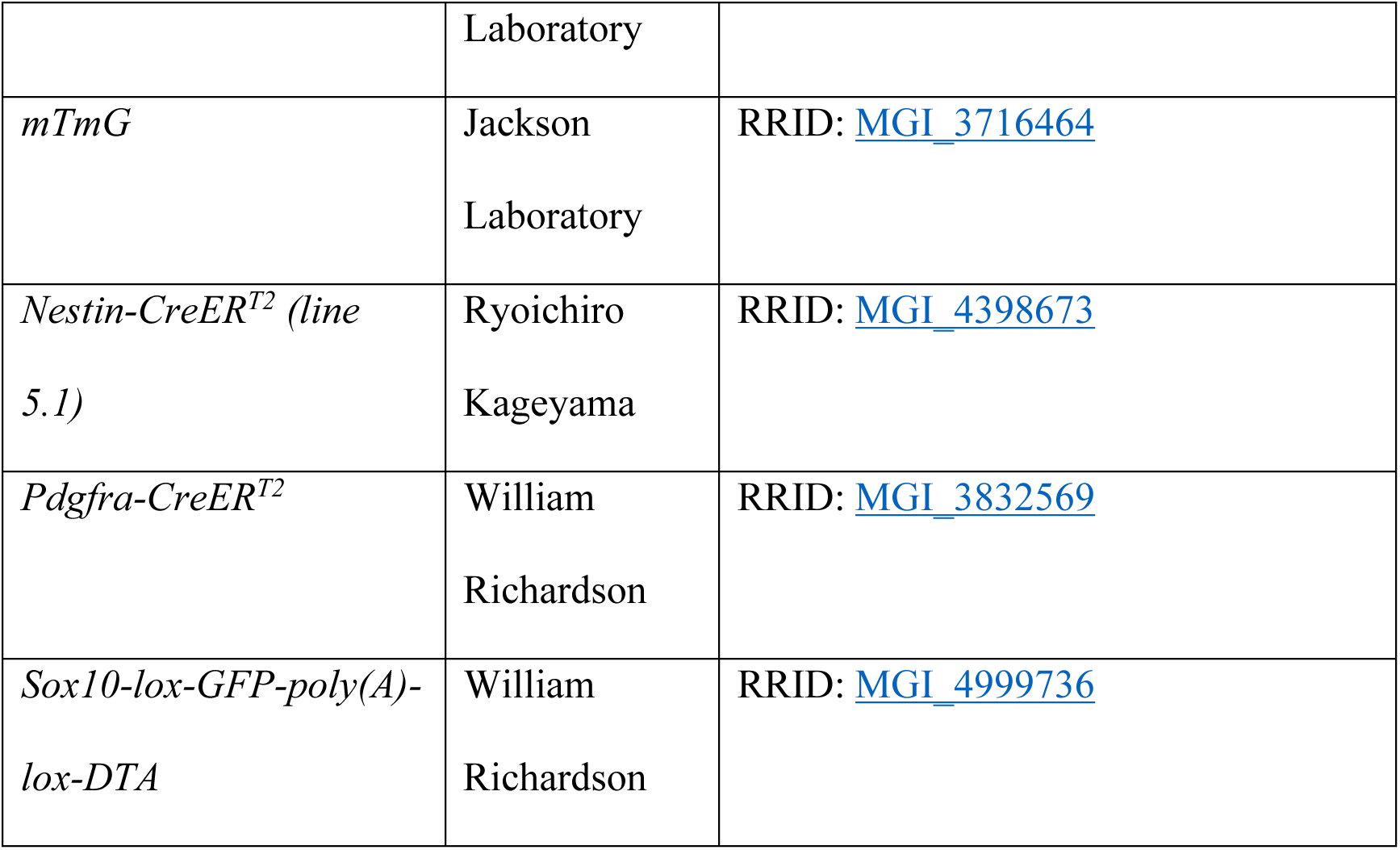
Key resources table

## Resource availability

### Lead contact

Further information and requests for resources and reagents should be directed to and will be fulfilled by the lead contact, Tobias D. Merson

### Materials availability

This study did not generate new unique reagents.

### Method details

#### Animals

Animal experiments were conducted in accordance with the National Health and Medical Research Council guidelines for the care and use of animals. All animal studies were approved by the animal ethics committee of the Florey Institute of Neuroscience and Mental Health (Parkville, VIC, Australia) and the animal ethics committee of Monash University (Clayton, VIC, Australia). *Pdgfrα-CreER^T2^*BAC transgenic mice (RRID: MGI_3832569) expressing CreER^T2^ under the regulation of the *Pdgfra* gene promoter, and *Sox10-DTA* transgenic mice (RRID: MGI_4999736) expressing a P1-derived artificial chromosome DNA construct containing the gene cassette *Sox10-lox-GFP-poly(A)-lox-DTA* driven by the *Sox10* promoter (Kessaris *et al*., 2006; Rivers *et al*., 2008). These two mouse lines were crossed to generate *Pdgfrα-CreER^T2+/+^: Sox10-DTA^+/−^* and *Pdgfrα-CreER^T2+/+^: Sox10-DTA^−/−^* breeders that were used to produce experimental cohorts comprising male and female offspring. To generate *Pdgfrα-CreER^T2+/−^: Sox10-DTA^+/−^: Ai14 tdTomato^+/−^* mice, we crossed *Pdgfrα-CreER^T2+/+^: Sox10-DTA^+/−^* mice with homozygous *Ai14 tdTomato^+/+^* mice (RRID: MGI_3809524), which were purchased from the Jackson Laboratory. We also generated *Nestin-CreER^T2+/–^; Pdgfrα-CreER^T2+/–^; Sox10-DTA^+/−^* transgenic mice for the combined ablation of both parenchymal OPCs and oligodendrogenic NPCs by crossing *Pdgfrα-CreER^T2+/+^: Sox10-DTA^+/−^* mice with *Nestin-CreER^T2^*^+/+^ *(line 5.1)* mice (RRID: MGI_4398673) (Imayoshi et al., 2006) generously provided by Ryoichiro Kageyama. We noted that a number of *Nestin^+^: Pdgfra^+^: DTA^+^* did not survive beyond weaning due to hydrocephalus. In surviving adult mice that were administered TAM and AraC (*n=*3 mice), we observed anatomical abnormalities consistent with hydrocephalus including expanded lateral ventricles (**Figure S9B**). Hydrocephalus was noted in *Nestin^+^: Pdgfra^+^: DTA^+^* mice as early as postnatal day 16. The ratio of *Nestin^+^: Pdgfra^+^: DTA^+^* versus *Nestin^+^: Pdgfra^+^: DTA^−^* mice surviving beyond weaning (P21) was 33.6% (36 out of 107 offspring) and gross hydrocephalus was evident in 39% of mice (14/36 mice) that survived post weaning. By contrast, no *Nestin^+^: Pdgfra^+^: DTA^−^* mice exhibited evidence of gross hydrocephalus (0/71 mice). The incidence of hydrocephalus in *Nestin^+^: Pdgfra^+^: DTA^+^* mice likely reflects TAM-independent recombination of the DTA allele due to leaky Cre activity driven by the *Nestin-CreER^T2^* allele during ontogeny thereby resulting in congenital apoptosis of a subset of neural crest cells (Plummer et al., 2017) which express both SOX10 and Nestin (Zhu et al., 2019). This possibility is supported by the observation that loss of neural crest cells during fetal development is documented to cause hydrocephalus (Guerra et al., 2015; Rodriguez et al., 2012). Finally, we generated *Nestin-CreER^T2+^: mTmG^+^* mice to evaluate the degree of TAM-independent recombination among adult neural progenitor cells by crossing *Nestin-CreER^T2^*^+/+^ *(line 5.1)* mice (RRID: MGI_4398673) with *mTmG^+/+^* mice (RRID: MGI_3716464).

#### TAM induction

Cre-mediated recombination was induced by oral gavage of TAM (Sigma) delivered at a dose of 300 mg/kg/d for 4 consecutive days, as described in previous studies (Rivers *et al*., 2008; Xing *et al*., 2014). TAM was prepared at 40 mg/mL in corn oil (Sigma). No toxicity due to TAM administration was observed in any cohort of mice.

#### EdU administration

To label cells that proliferated during the first 10 days following AraC withdrawal, 5-ethynyl-2ʹ-deoxyuridine (EdU; Life Technologies) was administered to mice in their drinking water at 0.1 mg/mL. EdU-supplemented drinking water was placed in light-proof water bottles and replaced every 3 days.

#### Preparation of AraC for intracisternal infusions

Cytosine-β-D-arabinofuranoside (AraC, Sigma) was prepared at a final concentration of 2% (w/v) in artificial CSF (aCSF, Tocris Bioscience). One hundred microliters of either 2% AraC or vehicle (aCSF) was injected into osmotic minipumps (Alzet Model 1007D, flow rate 0.5 µL/hour, Brain Infusion Kit III) using a 1 mL syringe attached to a blunt fill needle. The flow moderator was attached to a bespoke tubing assembly made by connecting PE-10 polyethylene tubing to the vinyl catheter tube provided with the Brain Infusion Kit III (Alzet). The flow moderator with attached tubing was slowly inserted into the filled osmotic minipump to create a complete pump assembly. The pumps were then transferred into 50 mL conical tubes containing sterile saline and placed in a 37^°^C water bath overnight to prime the pumps prior to surgical implantation.

#### Surgical implantation of osmotic minipumps

AraC or vehicle (aCSF) was infused into the CSF at the level of the cisterna magna via an osmotic minipump for a period of 6 days. Prior to anesthesia, mice received a subcutaneous injection of meloxicam (2 mg/kg, 0.25 mL/10 g body weight) in warm saline. Mice were then anaesthetized by isoflurane inhalation (4% induction, 2% maintenance). The head of the anesthetized mouse was fixed in a stereotaxic frame using a nose cone and ear bars. The position of the abdomen was lowered so that the neck was flexed at an angle of 30-45 degrees relative to horizontal and the body was placed on a thermostatically controlled heat pad to maintain body temperature. Eyes were moistened with water-based lubricant and the fur was cleared over the head and shoulders with an electric shaver. A sterile cotton tip soaked in 80% ethanol was used to swab and clean the surface of incision site, followed by 10% (w/v) povidone-iodine solution (Betadine). A midline skin incision was made using a sharp scalpel from a position just rostral of the external occipital protuberance to ∼1 cm cranial to the shoulders. The atlanto-occipital membrane was visualized after blunt dissection of the muscle layers to expose the position of the cisterna magna. Using a straight hemostat, a pocket was created by spreading the subcutaneous connective tissues apart and the osmotic minipump was inserted into the pocket. The cisterna magna was pierced superficially with a 30

G needle and the PE-10 tubing connected to the osmotic minipump was introduced into the hole before applying a small amount of superglue to fix the tubing in place. The position of the tubing was further anchored and fixed to the musculature using sutures. Bupivacaine (100 μL of 0.25% solution) was flushed over the musculature to provide rapid onset analgesia. The skin was then sutured and 10% (w/v) povidone-iodine solution was applied to the sutured skin. The animal was placed in a warm recovery box for monitoring until it regained consciousness and normal mobility. Animals were monitored daily throughout the experiment and were administered meloxicam (2 mg/kg, 0.25 mL/10 g body weight) in warm saline once daily for the first 2 days post-surgery. All mice were provided with powdered chow mixed with fresh water daily in a small dish that was easily accessed within the animal cage.

#### Surgical removal of osmotic minipumps

Infusion of AraC or vehicle (aCSF) was ceased after 6 days by removing the osmotic minipumps. Animals were anaesthetized as described above and the former skin incision site was reopened to gain access to the tubing connected to the osmotic minipump. The tubing was cut 2 mm from the glued/sutured musculature and the minipump was removed from the pocket. The tubing fixed to the musculature was left in place and the free end of the tubing was sealed with superglue. The incision was closed with sutures and the animal was placed in a warm recovery box for monitoring until it regained consciousness and normal mobility. Animals were monitored daily throughout the experiment. AraC-administered mice experienced a mild reduction in body weight during AraC infusion. If mice showed signs of greater than 10% weight loss, they were given powdered chow mixed with fresh water daily in a small dish that was easily accessed within the animal cage. If mice maintained greater than 15% weight loss for more than 72 hours, they were humanely euthanized. Following removal of the osmotic minipumps delivering AraC, mice returned to normal weight.

#### Generation of lentiviral vectors

The lentiviral vectors *LV-FUW-Nestin-NLS-HA-Dre* and *LV-FUW-EF1α-FREX-Myc/mKate2-f-mem* were used for fate-mapping of V-SVZ-derived NPCs. These vectors were constructed using standard molecular cloning techniques, including PCR using Phusion DNA polymerase (New England Biolabs), restriction enzyme digestion and Gibson assembly. To create these lentiviral vectors, the rat *Nestin* promoter sequence was amplified by PCR from plasmid DNA (Addgene plasmid #32401). The DNA encoding *NLS-HA-Dre* was amplified by PCR from plasmid DNA (Addgene plasmid #51272). The coding sequence for the rat *Nestin* second intron enhancer was amplified from rat genomic DNA. *EF1α* promoter sequence was amplified by PCR from plasmid DNA (Addgene plasmid #38770). The *FREX-Myc/mKate2-f-mem DNA* sequence was generated by DNA synthesis (Integrated DNA Technologies). The PCR products were cloned into the FUGW lentiviral vector backbone (Addgene plasmid #14883) in place of the GFP coding sequence by Gibson Assembly (New England Biolabs). Plasmid DNA was then extracted and purified using Plasmid Mini or Midi Kits (Qiagen). Sequences of the vector constructs were verified by DNA sequencing (Micromon, Monash University).

#### *In vitro* validation of lentiviral vectors

Plasmid DNA was transfected into HEK293T cells cultured at 37°C and 5% CO2 and analyzed 48 h post-transfection for fluorescence. The *pmKate2-f-mem* plasmid (Evrogen) served as a positive control for mKate2 fluorescence. HEK293T cells were plated in a 24 well plate and cultured in Dulbecco’s Modified Eagle’s Medium (DMEM, Gibco), supplemented with 10% fetal bovine serum (Invitrogen). At 80% confluency, cells were transfected with the plasmids using Lipofectamine 2000 Transfection Reagent (Invitrogen) according to the manufacturer’s instructions. The growth medium was replaced with fresh medium containing 100 U/mL penicillin and 100 mg/mL streptomycin (Gibco) 4 h post transfection. At 48 h after transfection, cells were post-fixed with 4% PFA/DPBS, processed for immunocytochemistry and imaged for mKate2 expression using a Zeiss LSM780 confocal microscope.

#### Lentivirus production

Lentiviruses were produced and packaged in HEK293T cells by the Vector and Genome Engineering Facility, Children’s Medical Research Institute. To determine viral titer, HEK293T cells were transduced with lentivirus. Genomic DNA was extracted from transduced cells and viral DNA copy number/cell was determined by Multiplex Taqman qPCR.

#### Intraventricular injection of lentiviral vectors

Mice were anaesthetized by isoflurane inhalation and positioned in a motorized stereotaxic frame as described above. A small incision was made in the scalp and the injection site of the lateral ventricle was marked using the following stereotaxic coordinates, relative to Bregma: anterior-posterior –0.22 mm, medial-lateral –1.00 mm, and dorsal-ventral –2.5 mm. A small hole was drilled into the skull to expose the brain surface. Five microliters of mixed lentiviral vectors were gently drawn up into a blunt-tipped 35G needle attached to a 10 µL NanoFil syringe (World Precision Instruments, WPI). The syringe was then placed into a microinjection pump (UltraMicroPump 4, WPI) attached to the stereotaxic frame and lowered slowly into the injection site of the lateral ventricle according to the dorsal-ventral coordinate. The microinjection pump controlled the infusion of 5 µL total volume of lentiviruses at a flow rate of 0.5 µL/min, after which the needle was left in place for 5 min to ensure complete diffusion of the viruses and avoid backflow.

#### Tissue processing and immunohistochemistry

Mice were deeply anesthetized with 100 mg/kg sodium pentobarbitone and then transcardially perfused with PBS, followed by 4% PFA/PBS. Brains, optic nerves and spinal cords were removed and post-fixed in 4% PFA/PBS for 2 h on ice, transferred to PBS overnight, cryopreserved in 20% sucrose/PBS overnight, followed by embedding in Tissue-Tek OCT compound (Sakura FineTek). The tissues were stored at –80°C until sectioned. Ten micron-thick coronal sections of the brain, spinal cord and optic nerve were cut on a Leica cryostat, collected onto Superfrost Plus slides (Menzel Glaser), and air dried for 1 h before storing at -80°C until stained. Cryosections were air dried, then blocked with PBS containing 0.3% Triton X-100, 10% normal donkey serum, and 10% BlokHen (Aves Labs) for 1 h at room temperature (RT). The sections were then incubated with primary antibodies at RT overnight, followed by 1 h incubation at RT with secondary antibodies.

For multiplex immunohistochemistry, some primary antibodies were incubated simultaneously. The following primary antibodies were used: rabbit anti-ALDH1L1 (1:1000, Abcam, Cat. ab87117), mouse anti-CC1 (1:100, Calbiochem, Cat. OP80), rat anti-CD16/CD32 (1:100, BD Biosciences, Cat. 553142), rabbit anti-CD206 (1:200, Abcam, Cat. ab64693), goat anti-DCX (1:100; Santa Cruz Biotechnology, Cat. sc-8066), mouse anti-FoxJ1 (1:200, eBioscience, Cat. 14-9965-82), mouse anti-GFAP (1:500, Millipore, Cat. MAB360), chicken anti-GFP (1:2000, Aves Labs, Cat. GFP-1020), rabbit anti-GPR17 (1:800, Cayman, Cat. 10136), mouse anti-HA-tag (1:500, Sigma, Cat. H9658), rabbit anti-Iba1 (1:200, Wako, Cat. 019-19741), goat anti-Iba1 (1:500, Abcam, Cat. ab5076), rabbit anti-Laminin-1 (1:400, Sigma, Cat. L9393), mouse anti-Myc-tag (1:500, Sigma, Cat. 05-419), mouse anti-Nestin (1:100, Millipore, Cat. MAB353), mouse anti-NeuN (1:100, Millipore, Cat. MAB377), rabbit anti-NG2 (1:200, Millipore, Cat. AB5320), goat anti-PDGFRA (1:150, R&D Systems, Cat. AF1062), rat anti-PDGFRA (1:150, BD Biosciences, Cat. 558774), goat anti-PDGFRB (1:200, R&D Systems, Cat. AF1042), mouse anti-α-SMA (1:500, Abcam, Cat. ab7817), goat anti-SOX10 (1:100, R&D Systems, Cat. AF2864), and rabbit anti-TagRFP for mKate2 labelling (1:500, Kerafast, Cat. EMU113). To label GFP^+^ tdTomato^+^ brain sections using primary antibodies against Nestin, PDGFRA, and NG2, as well as EdU and Hoechst, inactivation of GFP and tdTomato was performed by firstly treating the brain sections with 3% H_2_O_2_ and 20 mM HCl in PBS for 1 h at RT with light illumination (Lin et al., 2016). The slides were then washed three times with PBS, incubated with blocking buffer and processed for the immunostaining as described above.

Secondary antibodies raised in donkey and conjugated to Alexa Fluor 488, FITC, TRITC, Alexa Fluor 594 or Alexa Fluor 647 were purchased from Jackson ImmunoResearch or Invitrogen and used at 1:200 dilution. Sections incubated with biotinylated rat anti-PECAM1/CD31 antibody (1:200, BD Biosciences, Cat. 553371) were rinsed and further incubated with streptavidin-Brilliant Violet 480 (1:200; BD Biosciences) for 30 min. Some slides stained without the fluorophore Brilliant Violet 480 were also counterstained with Hoechst 33342 (1 µg/mL, Invitrogen).

For myelin analysis, slides were stained with Black-Gold II (Biosensis) according to the manufacturer’s instructions. To detect EdU incorporation in proliferating cells, sections were first processed for immunohistochemistry as above, followed by EdU detection using the Click-iT EdU Alexa Fluor 647 Imaging Kit (Life Technologies) as per the manufacturer’s instructions. Sections were coverslipped with Mowiol mounting medium and subjected to fluorescence and confocal microscopic analysis.

#### Fluorescence imaging and image analysis

Stained 10 µm thick coronal sections were imaged by laser scanning confocal microscopy (Zeiss LSM510-META or Zeiss LSM780), which was used to detect up to four fluorophores by laser excitation at 405, 488, 561 and 633 nm wavelengths. For five-color imaging such as brain sections stained with Brilliant Violet 480 or Hoechst, as well as Alexa Fluor 488, TRITC, Alexa Fluor 594 and Alexa Fluor 647, linear unmixing was performed during acquisition (online fingerprinting). Tile scanning was performed at a magnification of 10x or 20x for the cellular analysis of the entire brain sections of transgenic mouse lines. Confocal images were then imported into ImageJ for quantification of cellular density in the regions of interest.

For global analysis of cellular distribution in the entire mouse brain across Bregma levels, 2-3 random sections were scanned from the stained slides on an Olympus VS120 Virtual Slide Microscope with a 20x objective at Monash Histology Platform, Monash University. The resulting images have a pixel resolution of 0.65 µm/pixel. Double- or triple-positive cells from these images were automatically counted using ImageJ scripts created by Monash Micro Imaging, Monash University. Results obtained from automatic cell quantification were also validated by manual counts. For the analysis of Laminin-1/PDGFRA, NG2/PDGFRA or PDGFRB/PDGFRA colocalization, confocal images were acquired using a 63X objective to generate z-stacks. Complete image stack deconvolution was performed with the “Iterative Deconvolve 3D” plugin in ImageJ. Colocalization analysis was performed by using the “Colocalization threshold” plugin in ImageJ to automatically determine a detection threshold for each channel to avoid subjective bias. The extent of within-pixel fluorescent signal colocalization as indicated by Pearson’s correlation coefficient was calculated in each optical slice and then flattened into a maximal z-projection to reveal colocalized pixels across the entire image thickness. All analyses were performed in a blinded fashion.

For morphological analysis of microglia and astrocytes, Z-stacks of 1-4 cells were taken from the cortical region of each of the brain sections under a 40x objective. Microglia soma and branching measures were visualized using IBA1 immunofluorescence, whereas those for astrocytes were assessed with GFAP. Z-stacked images were converted to maximum intensity projections using Fiji/ImageJ and these images were background subtracted, contrast-enhanced to ensure full arborization could be detected and a local threshold was applied to the image. Microglial and astrocytic soma areas were calculated using the ‘Measure’ command and branch features (number of primary or secondary processes; maximum length of primary process) were manually counted and measured in Fiji/ImageJ. 18-22 astrocytes and 18-39 microglia per group were analyzed in a blinded fashion.

#### Bright-field imaging and image analysis

For the sections stained with Black-Gold II, 2-3 representative images were taken using a 10x objective on a Zeiss Axioplan upright fluorescent microscope and captured with an Axioplan HRc camera (Carl Zeiss) using the Axiovision 7.2 imaging software. All images were taken with the same exposure time. For quantification of myelin intensity, images from the sections stained with Black-Gold II were converted to grayscale in ImageJ and automated measurements of myelin intensity were taken using the measurement function of ImageJ to record the mean gray value within the regions of interest. All analyses were performed in a blinded fashion.

#### Statistical analyses

All statistical analyses were performed using the GraphPad Prism software. Statistical significance was determined using an unpaired, two-tailed Student’s *t*-test or by two-way ANOVA with Bonferroni’s, Tukey’s or Sidak’s multiple-comparison tests. Statistical significance was defined as *p*<0.05. Quantitative data are reported as mean ± SEM.

## Acknowledgments

This work was supported by funding from the Australian Research Council Special Research Initiative - Stem Cells Australia (T.D.M., T.J.K.) and was supported (in part) by the Intramural Research Program of the NIMH. Y.L.X received support from The CASS Foundation (8545). T.D.M was the recipient of a Future Fellowship from the Australian Research Council (FT150100207) and received philanthropic support from Metal Manufactures Ltd. We thank Ryoichiro Kageyama (RIKEN Center for Brain Science, Japan) for generously providing the *Nestin-CreER^T2^ (line 5.1)* mice. The authors would also like to acknowledge the use of facilities within the Monash Animal Research Platform, Monash Micro Imaging Platform, Monash Histology Platform, and Micromon (Monash University).

## Author contributions

Y.L.X., W.D.R., T.J.K. and T.D.M. designed the research. Y.L.X., Y.O., J.P., B.H.A.C., K.M., S.M. and T.D.M. performed research. Y.L.X. and T.D.M. analyzed and interpreted data and wrote the manuscript.

## Corresponding author and lead contact

Tobias D. Merson

## Data availability statement

The authors declare that the data supporting the findings of this study are available within the paper and its supplementary information files.

