## Supplemental Figure Legends for "High-Efficiency Pharmacogenetic Ablation of Oligodendrocyte Progenitor Cells in the Adult Mouse CNS"

**Figure S1. DTA-mediated ablation of OPCs induced rapid cell regeneration.**

**A,B**, All residual GFP$^+$ cells in *Pdgfrα$^+$: DTA$^+$* mice examined 4 days after TAM administration expressed Sox10 and CC1 (yellow arrowheads). A subpopulation of Sox10$^+$ oligodendroglia did not express GFP (**A**, white arrowheads) suggesting that the *Sox10-lox-GFP-STOP-lox-DTA* allele is not transcriptionally active in all Sox10$^+$ oligodendroglia. **C**, EdU labeling of PDGFRA$^+$ cells in *Pdgfrα$^+$: DTA$^+$* mice at 8 days post TAM administration. **D**, Densities of PDGFRA$^+$ OPCs and their subpopulations expressing GFP and/or EdU in the corpus callosum of *Pdgfrα$^+$: DTA$^+$* mice at 4, 8 and 10 days after the final tamoxifen administration (n=4 mice per group). **E**, Densities of PDGFRA$^+$ OPCs in the corpus callosum of wild-type mice infused with vehicle (aCSF) or AraC for 6 days before analysis at 0 or 10 dppr. Data represent mean ± SEM. Statistics: Two-way ANOVA with Tukey's multiple comparison tests, *$p$<0.05, ****$p$<0.0001 Scale bars, 50 µm (**A**) and 40 µm (**B,C**).

**Figure S2. Intracisternal infusion of AraC following TAM-induced DTA expression in OPCs results in highly efficient OPC ablation throughout the brain.**

**A-C**, Immunohistochemistry against PDGFRA, PECAM-1, Laminin-1, NG2 and PDGFRB in coronal sections reveals vascular-associated PDGFRA$^+$ cells are perivascular fibroblasts-like cells surrounding PECAM-1$^+$ endothelial cells. Colocalization analysis of PDGFRA with Laminin-1, a key vascular basement membrane component (**A**) and pericyte markers NG2 and PDGFRB (**B,C**) in a TAM + vehicle administered *Pdgfrα $^+$: DTA$^-$* mouse (**A,B**) and a TAM + AraC administered *Pdgfrα$^+$: DTA$^+$* mouse (**C**). Frequency scatterplots of fluorescence intensity in the red and green channels and photomicrographs showing colocalized red and green pixels (bottom right panels, in white) revealed a high degree of colocalization between PDGFRA and Laminin-1 (**A',A"**) whereas the degree of colocalization between PDGFRA and NG2 (**B',B"**) or PDGFRA and PDGFRB (**C',C"**) was low. **D**, Plot of Pearson's r coefficient of colocalization between PDGFRA and other antigens. **E,** Immunohistochemical detection of PDGFRA in serial coronal brain sections collected from non-ablated controls (upper row) and OPC-ablated mice (lower row) at the end of AraC or vehicle infusion (0 dppr). The non-ablated control group comprised *Pdgfrα*$^{+}$*: tdT*$^{+}$*: DTA*$^{-}$ mice administered TAM + vehicle. The OPC-ablated group consisted of *Pdgfrα*$^{+}$*: tdT*$^{+}$*: DTA*$^{+}$ mice administered TAM + AraC. The approximate position along the rostro-caudal axis for each coronal section is indicated below the photomicrographs. **F**, High magnification of the boxed region from (E) reveals that remnant PDGFRA$^{+}$ cells in OPC-deficient mice do not exhibit typical OPC morphology and are localized around PECAM-1$^{+}$ endothelial cells within the brain vasculature. **G**, Densities of tdTomato$^{+}$ PDGFRA$^{+}$ perivascular cells in coronal sections of non-ablated and OPC-ablated mice assessed at various rostrocaudal positions at 0 dppr. Data represent mean ± SEM. Statistical analysis: one-way (D) or two-way (G) ANOVA with Bonferroni's *post hoc* analysis, **p*<0.05, ***p*<0.01, *****p*<0.0001. Scale bars, 20 μm (**A**) and 30 μm (**B,C**), 900 μm (**E**) and 100 μm (**F**).

**Figure S3. PDGFRA-expressing cells remain depleted in the cerebrum for at least 10 days post infusion of AraC but are regenerated within 20 days post infusion of AraC.** **A,** Immunohistochemistry against PDGFRA on coronal brain sections of a TAM + AraC-administered *Pdgfrα*$^{+}$*: DTA*$^{+}$ mouse at 10 dppr reveals that ramified PDGFRA$^{+}$ cells remained depleted in the cerebrum (CH) but had started to repopulate more caudoventral regions of the brain such as brain stem (BS). The approximate rostro-caudal position relative to Bregma is indicated below each photomicrograph. **B,** Immunohistochemistry against

PDGFRA on coronal brain sections of a TAM + AraC-administered *Pdgfrα⁺: DTA⁺* mouse examined at 20 dppr reveals significant repopulation of both the cerebrum and brain stem with PDGFRA+ cells. The approximate rostro-caudal position relative to Bregma is indicated below each photomicrograph. **C**, *Pdgfrα⁺: DTA⁺* mice administered TAM + AraC and assessed at 10 dppr revealed that unlike dorsal regions of the cerebrum where OPCs remained depleted, PDGFRA-expressing cells in the caudoventral brain such as brain stem had started to regenerate. By 20 dppr, only caudodorsal regions of OPC-ablated mice were not fully repopulated by $PDGFRA^+$ cells when compared to non-ablated TAM + vehicle (0 dppr) mice, two-way ANOVA with Bonferroni's *post hoc* analysis, $^{****}p<0.0001$ (overall effect). **D**, Densities of $PDGFRA^+$ only and $PDGFRA^+$ $tdTomato^+$ cells in the brain at 0 and 20 dppr. BS, brain stem; CH, cerebrum. Scale bar, 1 mm.

**Figure S4. Consequences of pharmacogenetic ablation on cells at intermediate stages of OL differentiation and phenotypic characterization of newly-generated $PDGFRA^+$ cells in the corpus callosum proximal to the V-SVZ.**

**A,** Coronal sections of the rostral corpus callosum of a non-ablated control (*Pdgfrα⁺: DTA⁻*, TAM + vehicle) and OPC-ablated mouse (*Pdgfrα⁺: DTA⁺*, TAM + AraC) collected at 0 and 20 dppr and immunostained with antibodies against PDGFRA and GPR17. **B,** Densities of cells expressing PDGFRA and/or GPR17 in the rostral corpus callosum at the assessed time-points. Early OPCs ($PDGFRA^+$ $GPR17^-$ and $PDGFRA^+$ $GPR17^+$) and intermediate pre-OLs ($PDGFRA^-$ $GPR17^+$) were completely ablated at both 0 and 10 dppr (n=4 mice per group, mean ± SEM). **C,** Coronal sections of the rostral corpus callosum of an OPC-deficient mouse assessed at 20 dppr and labelled with antibodies against GFP, PDGFRA and NG2. $PDGFRA^+$ $NG2^+$ cells expressing the *Sox10-GFP* transgene (white arrows) that exhibited typical OPC-like morphology were identified proximal to the V-SVZ. CC, corpus callosum; Ctx, cerebral cortex; LV, lateral ventricle. Scale bars, 75 µm (**A**)**,** 100 µm (**C**) and 40 µm (**C'**).

**Figure S5. Phenotypic analysis of GFAP$^+$ astrocytes in the brain following OPC ablation.**

**A,** Density of GFAP$^+$ astrocytes in the brain was similar in OPC-ablated versus non-ablated controls (n=3 mice per group, mean ± SEM). Overall, the density of GFAP$^+$ astrocytes declined with time post-infusion (effect of time: *p*=0.0192, two-way ANOVA with Sidak's multiple comparisons test). **B,** Soma area of GFAP$^+$ astrocytes was similar in OPC-ablated versus non-ablated controls and increased in both groups with time post-infusion (effect of time: *p*=0.0007, two-way ANOVA with Sidak's multiple comparisons test). **C,D,** The number of primary and secondary processes did not differ between groups but both groups exhibited a small increase in process number with time post-infusion (effect of time: p=0.0011 (C), p=0.0363 (D), two-way ANOVA with Sidak's multiple comparisons test). **E,** The maximum length of primary processes was similar between OPC-ablated and non-ablated controls, but there was an overall effect of time post-infusion (p=0.0007, two-way ANOVA with Sidak's multiple comparisons test).

**Figure S6. Phenotypic analysis of Iba1$^+$ microglia in the brain following OPC ablation.**

**A,** Modest changes in mean soma size of Iba1$^+$ microglia in OPC-ablated and non-ablated controls with time post-infusion. Two-way ANOVA with Sidak's multiple comparisons test revealed an overall effect of time (*p*=0.0052). **B,** Reduction in the number of primary processes of microglia in OPC-ablated mice at 10 dppr. Two-way ANOVA with Sidak's multiple comparisons test revealed an overall effect of time (*p*=0.0479) and treatment (*p*=0.0149) and an interaction between these variables (*p*<0.0001). **C,** Dynamic changes in the number of secondary processes of microglia in OPC-ablated and non-ablated controls.

Two-way ANOVA with Sidak's multiple comparisons test revealed an overall effect of time ($p$=0.0005) and treatment ($p$=0.043) and an interaction between these variables ($p$<0.0001). **D,** Reduction in the maximum length of primary processes of microglia in OPC-ablated mice at 10 and 20 dppr. Two-way ANOVA with Sidak's multiple comparisons test revealed an overall effect of time ($p$=0.0092) and treatment ($p$=0.0011). *$p$<0.05, ****$p$<0.0001. **E,** Iba1 and CD16/CD32 immunohistochemistry performed on sections of the cerebral cortex of OPC-ablated and non-ablated controls at 0, 10 or 20 dppr. Scale bar, 110 µm. **F,** Quantification of Iba1$^+$ cell density according to presence or absence of either CD16/CD32 or CD206 at 0, 10 and 20 dppr in OPC-ablated and non-ablated controls.

**Figure S7. Dynamics of ablation and regeneration of PDGFRA$^+$ cells throughout the central nervous system.**

**A**, Coronal section of the rostral forebrain of a *Pdgfrα$^+$: tdT$^+$: DTA$^-$* mouse administered TAM + vehicle at 20 dppr reveals that tdTomato was expressed in recombined PDGFRA$^+$ OPCs and in oligodendroglia that have differentiated from fate-mapped OPCs, as well as in some PDGFRA$^+$ perivascular fibroblast-like cells. **B**, By contrast, *Pdgfrα$^+$: tdT$^+$: DTA$^+$*, TAM + AraC mice at 20 dppr exhibited repopulation of PDGFRA$^+$ cells, with the highest cell density in the corpus callosum proximal to the V-SVZ and lower densities in the septum, caudate putamen and cerebral cortex. Interestingly, tdTomato expression was not observed in these regenerated cells and it was only expressed in non-ablated perivascular fibroblast-like cells. **C**, Densities of PDGFRA$^+$ cell subpopulations in the corpus callosum and cerebral cortex of *Pdgfrα$^+$: tdT$^+$: DTA$^-$* and *Pdgfrα$^+$: tdT$^+$: DTA$^+$* mice administered TAM + vehicle or TAM + AraC and assessed at either 0 or 20 dppr (n=4 mice per group, mean ± SEM). (****$p$<0.0001, two-way ANOVA with Bonferroni's *post hoc* analysis). **D**, In the spinal cord and optic nerves of TAM + vehicle administered *Pdgfrα$^+$: tdT$^+$: DTA$^-$* mice, tdTomato was expressed in the majority of PDGFRA$^+$ OPCs and in oligodendroglia that had differentiated from fate-mapped OPCs. **E**, Fate-mapped OPCs were ablated in the coronal spinal cord and optic nerve of TAM + AraC administered *Pdgfrα*$^+$*: tdT*$^+$*: DTA*$^+$ mice and assessed at 0 dppr. By 20 dppr, PDGFRA$^+$ cells had regenerated with a mixture of tdTomato-positive and tdTomato-negative cells. **F,G**, Densities of PDGFRA$^+$ cells in the coronal spinal cord **(F)** and optic nerve (**G**) of *Pdgfrα*$^+$*: tdT*$^+$*: DTA*$^-$ and *Pdgfrα*$^+$*: tdT*$^+$*: DTA*$^+$ mice administered TAM + vehicle or TAM + AraC and assessed at either 0 or 20 dppr (n=3-4 mice per group, mean ± SEM), (*$p$<0.05, **$p$<0.01, ***$p$<0.001, ***$p$<0.0001, two-way ANOVA with Tukey's *post hoc* analysis). CC, corpus callosum; CPu, caudate putamen; CSp, coronal spinal cord; Ctx, cerebral cortex; LV, lateral ventricle; OpN, optic nerve. Scale bars, 700 µm (**A,B**), 250 µm (Csp) and 100 µm (OpN) (**D,E**).

**Figure S8. Lentiviral approach for the fate-mapping of V-SVZ-derived NPCs.**

**A**, Schematic map for the construction of the *FUW-EF1α-FREX-Myc/mKate2-f-mem* plasmid. **B**, Schematic map for the construction of the *FUW-Nestin-NLS-HA-Dre* plasmid. **C**, Upon Dre-mediated recombination, the coding sequence of Myc/mKate2-f-mem is flipped into the correct sense orientation resulting in the expression of Myc-tagged, membrane-targeted mKate2 under the control of the human *EF1α* promoter. **D**, *In vitro* validation of plasmids showing HEK293T cells were transfected with commercial *pmKate2-f-mem* (Evrogen) as a positive control or *pNestin-NLS-HA-Dre* and *pEF1α-FREX-Myc/mKate2-f-mem*. Immunostaining shows mKate2 expression in the transfected HEK293T cells. Nuclei were stained with Hoechst. **E**, *In vivo* validation of lentiviral vectors showing the co-expression of mKate2 and Myc-tag or HA-tag in the V-SVZ wall of the lateral ventricle at 2 weeks after intraventricular injection of lentiviruses into wild type *C57BL/6J* mice. **F,G**, mKate2 expression was also observed in transduced multi-ciliated ependymal cells immunoreactive for α-SMA (**F**, yellow arrowed) or FoxJ1 (**G**) in addition to NPCs (**F**, white arrowed) in the wild-type V-SVZ. LV, lateral ventricle. **H,I,** Lentiviral-transduced cells expressing Myc-tag in the V-SVZ of transgenic mice at 20 days post AraC infusion. Membrane-targeted Myc-tag expression was observed in the V-SVZ, including D-SVZ (**H**) and DL-SVZ (**I**). Scale bars, Scale bars, 180 μm (**D**), 100 μm (**E**), 24 μm (**F**), 12 μm (**G**), 40 μm (**H**) and 85 μm (**I**).

**Figure S9. Repopulation of PDGFRA$^+$ cells is blocked by co-ablation of parenchymal OPCs and oligodendrogenic NPCs**.

**A,** Immunohistochemistry against PDGFRA on coronal brain sections of mice depleted of both parenchymal OPCs and oligodendrogenic NPCs (TAM + AraC-administered *Nestin$^+$:Pdgfra$^+$:DTA$^+$* mice), as compared to non-ablated controls (TAM + vehicle administered *Nestin$^+$:Pdgfra$^+$:DTA$^-$* mice). Ramified PDGFRA$^+$ cells remained depleted in the dorsal cerebrum (CH) of TAM + AraC-administered *Nestin$^+$:Pdgfra$^+$:DTA$^+$* mice assessed at 20 dppr but had started to repopulate more caudoventral regions of the brain particularly the brain stem (BS). The enlarged ventricles of *Nestin$^+$:Pdgfra$^+$:DTA$^+$* mice are indicated by blue dashed line. **B,** Plot of lateral ventricle area in mice depleted of both parenchymal OPCs and oligodendrogenic NPCs versus non-ablated control mice (n=3-4 mice per group, mean ± SEM) (***$p$<0.0002, two-way ANOVA with Bonferroni's *post hoc* analysis.). **C,D,** Densities of DCX$^+$ cells in the lateral wall of the lateral ventricle (**c**) and in the dorsolateral corner of the lateral ventricles (**D**) in mice depleted of both parenchymal OPCs and oligodendrogenic NPCs as compared to non-ablated controls. **E,** Immunohistochemistry against PDGFRA, Laminin-1 and GFP on a coronal section of the cerebral cortex of a *Pdgfrα$^+$: tdT$^+$: DTA$^+$* mouse administered TAM + AraC and collected at 20 dppr provides no evidence that newly-generated PDGFRA$^+$ cells emerge from the meningeal membrane. **E'**, High magnification of the boxed region shown in (**E**) reveals no evidence of PDGFRA$^{+}$ cell regeneration near the meninges. **E''**, By contrast, a few PDGFRA+ cells were observed in the deeper cortical regions. **F**, Representative coronal section from the cerebral cortex of a *Pdgfrα$^{+}$: tdT$^{+}$: DTA$^{+}$* mouse administered TAM + AraC and collected at 20 dppr was immunostained with antibodies against PDGFRA, NG2 and Nestin, as well as Hoechst and EdU after fluorophore inactivation. The image demonstrates the absence of newly-generated PDGFRA$^{+}$ NG2$^{+}$ EdU$^{-}$ cells near the meninges. **G,** Extent of TAM-independent recombination among dividing cells (PCNA$^{+}$) and neuroblasts (DCX$^{+}$) in the V-SVZ of untreated adult *Nestin-CreER$^{T2+}$: mTmG$^{+}$* mice. In this line, the tdTomato reporter protein is constitutively expressed under the regulatory control of the *Rosa26* locus unless and until the *mTmG* allele undergoes Cre-mediated recombination, which results in excision of the tdTomato CDS and places the GFP CDS under control of the *Rosa26* locus. *Nestin-CreER$^{T2+}$: mTmG$^{+}$* mice (n=5) were sacrificed at 12 weeks of age and coronal sections were immunolabeled for GFP, PCNA and DCX. Only 0.95 ± 0.40% of DCX$^{+}$ cells in the V-SVZ co-expressed GFP, and 0.10 ± 0.23% of PCNA$^{+}$ cells in the V-SVZ co-expressed GFP, indicating that TAM-independent recombination occurs in <1% of NPCs. Scale bars, 100 μm (**E**) and 40 μm (**F**).
